# Remarkable recent changes in genetic diversity of the avirulence gene *AvrStb6* in global populations of the wheat pathogen *Zymoseptoria tritici*

**DOI:** 10.1101/2020.09.18.303370

**Authors:** Christopher Stephens, Fatih Ölmez, Hannah Blyth, Megan McDonald, Anuradha Bansal, Emine Burcu Turgay, Florian Hahn, Cyrille Saintenac, Vladimir Nekrasov, Peter Solomon, Andrew Milgate, Bart Fraaije, Jason Rudd, Kostya Kanyuka

## Abstract

Septoria tritici blotch (STB), caused by the fungus *Zymoseptoria tritici*, is one of the most economically important diseases of wheat. Recently, both factors of a gene-for-gene interaction between *Z. tritici* and wheat, the wheat receptor-like kinase Stb6 and the *Z. tritici* secreted effector protein AvrStb6, have been identified. Previous analyses revealed a high diversity of *AvrStb6* alleles present in historic *Z. tritici* isolate collections, with up to ~ 18% of analysed isolates possessing the avirulence isoform of AvrStb6 identical to that originally identified in the reference isolate IPO323. With *Stb6* present in many commercial wheat cultivars globally, we aimed to assess potential changes in *AvrStb6* genetic diversity and the incidence of alleles allowing evasion of *Stb6*-mediated resistance in more recent *Z. tritici* populations. Here we show, using targeted re-sequencing of *AvrStb6,* that this gene is universally present in field isolates sampled from major wheat-growing regions of the world between 2013–2017. However, in contrast to the data from studies of historic isolates, our study revealed a complete absence of the originally described avirulence isoform of AvrStb6 amongst modern *Z. tritici* isolates. Moreover, a remarkably small number of alleles, each encoding AvrStb6 protein isoforms conditioning virulence on *Stb6-*containing wheat, were found to predominate among modern *Z. tritici* isolates. A single virulence isoform of AvrStb6 was found to be particularly abundant throughout the global population. These findings indicate that, despite the ability of *Z. tritici* to sexually reproduce on resistant hosts, *AvrStb6* avirulence alleles tend to be eliminated in subsequent populations.

## INTRODUCTION

The interactions between plant pathogens and their hosts during infection are highly complex and evolutionarily dynamic. Effectors, molecules including proteins that are produced and secreted by pathogens during infection, constitute a vital part of the repertoire of mechanisms utilised in the successful infection of plant hosts (Lo Presti *et al.*, 2015).

Effectors function by altering the metabolism of the host plant to facilitate infection, or by suppressing plant immune responses to infection (Djamei *et al.*, 2011; Marshall *et al.*, 2011; Kleemann *et al.*, 2012). However, plants possess a sophisticated innate immune system whose central players are cell surface and intracellular immune receptors including disease resistance (*R*) genes, which are capable of detecting pathogen effectors and initiating an immune response (Jones and Dangl, 2006; Cook *et al.*, 2015; Kanyuka and Rudd, 2019).

Recognition of a pathogen effector protein by a plant immune receptor thereby introduces an evolutionary pressure on the pathogen to mutate or lose the recognised effector (otherwise known as an avirulence or Avr factor) entirely from its genome. Avr factors may be lost through frameshift or nonsense mutations (Luderer *et al.*, 2002), transposon insertion (Zhang *et al.*, 2015), or repeat-induced point mutations (RIPs) (Rouxel *et al.*, 2011). Indeed, effectors are often located in genomic regions rich in transposon activity which drives effector diversity (Dong *et al.*, 2015). Suppression of Avr factor expression, through mutations in the promoter, histone modification (Soyer *et al.*, 2014) or post-transcriptional silencing (Qutob *et al.*, 2013) may also restore pathogen virulence. In some cases, point mutations in the *Avr* gene sequence may allow evasion of recognition whilst maintaining protein function (Blondeau *et al.*, 2015; Plissonneau *et al.*, 2017). This may happen in cases where the Avr factor is important for pathogen fitness. Once mutations in *Avr* genes have occurred that allow evasion of detection by immune receptors, they often spread rapidly through the pathogen population particularly when the cognate disease resistance gene is widely used (Cowger *et al.*, 2000; Hovmøller and Justesen, 2007).

*Zymoseptoria tritici*, the causal agent of Septoria tritici blotch (STB), is one of the most economically important fungal pathogens of wheat, with fungicide control costs exceeding €1 billion per year in Europe alone (Torriani *et al.*, 2015). *Z. tritici* secretes an array of putative effectors during the infection cycle, among which is the avirulence factor AvrStb6 (Zhong *et al.*, 2017; Kema *et al.*, 2018), recognised in a gene-for-gene manner by the cognate cell surface immune receptor Stb6 (Brading *et al.*, 2002; Saintenac *et al.*, 2018). *AvrStb6* has been identified in the genomes of the isolates IPO323 and 1E4 collected from the Netherlands and Switzerland in 1981 and 1999, respectively, and recently cloned (Zhong *et al.*, 2017; Kema *et al.*, 2018). The *Stb6* gene has also been recently cloned from wheat landrace Chinese Spring and the UK cultivar (cv.) Cadenza (Saintenac *et al.*, 2018). *AvrStb6* and *Stb6* encode a small, cysteine-rich secreted protein and a wall-associated receptor-like kinase protein, respectively. A functional haplotype of *Stb6* conferring resistance to *Z. tritici* IPO323 is found in over half of commercial cultivars in Europe (Saintenac *et al.*, 2018) and may have been present in wheat populations globally since early in agricultural history (Chartrain *et al.*, 2005). Consequently, pressure on *Z. tritici* to lose *AvrStb6* variants conditioning avirulence on *Stb6* containing wheat must be considerable.

There are two previous studies, in which sequencing of *AvrStb6* from historic populations of *Z. tritici* has been carried out. In one study, a global population of *Z. tritici* collected between 1990–2001 (Zhan *et al.*, 2005; Brunner and McDonald, 2018) was analysed, whilst the second focussed on a largely French population collected in 2009–2010 (Zhong *et al.*, 2017). These studies found a high diversity of *AvrStb6* alleles, and evidence of positive selection driven by point mutations and recombination. The region of *Z. tritici* chromosome 5 in which *AvrStb6* is located was found to be highly dynamic, with extensive transposon activity contributing to *AvrStb6* polymorphism (Sánchez-Vallet *et al.*, 2018).

Whilst the *AvrStb6* allele distribution in these historic populations have been well characterised, the *AvrStb6* allele diversity in more modern *Z. tritici* populations is unknown. Also unknown is the prevalence of *Z. tritici* isolates which possess *AvrStb6* alleles capable of evading *Stb6*-mediated defence. The specific polymorphisms that drive the change from avirulence to virulence phenotype in the AvrStb6 protein have not yet been determined, although changes at the two amino acid residues (positions 41 and 43) in the AvrStb6 protein have been suggested as being critical for the pathogenicity on wheat cultivars carrying *Stb6* (Kema *et al.*, 2018).

In this study, we re-sequenced the *AvrStb6* gene from recent field populations of *Z. tritici* isolates collected between 2013–2017 therefore providing a global snapshot of *AvrStb6* allele diversity at this time. We show a notable decrease in the allele diversity of *AvrStb6*, compared to previous studies, including a recent worldwide shift towards allele encoding *Stb6* resistance-breaking isoforms. Interestingly, one particular virulence isoform (I02) was found to be the most abundant in several regions of the world examined. This study therefore provides a rare insight into temporal changes in pathogen effector diversity in response to the host-imposed pressures and highlights the speed with which these changes can occur when a single cognate disease resistance gene is deployed extensively.

## RESULTS

### *AvrStb6* allele analysis

Primers flanking the *AvrStb6* gene were designed and used for PCR amplification of this gene from a collection of 383 *Z. tritici* isolates sampled from different field grown bread wheat (*Triticum aestivum*) cultivars in ten countries on five continents. PCR products of expected size were obtained from all analysed samples indicating *AvrStb6* is present in all isolates in the collection. Sequencing of the PCR products revealed a total of 52 *AvrStb6* alleles (denoted from A01 through to A52; Table S1) coding for 44 protein isoforms (denoted from I01 through to I44; Figure 1). *AvrStb6* alleles and the corresponding protein isoforms from the reference isolate IPO323 and another well-studied isolate IPO88004, which are avirulent or virulent on *Stb6-*containing wheat, respectively, were named as A01/I01 and A52/I44.

**Figure 1.**
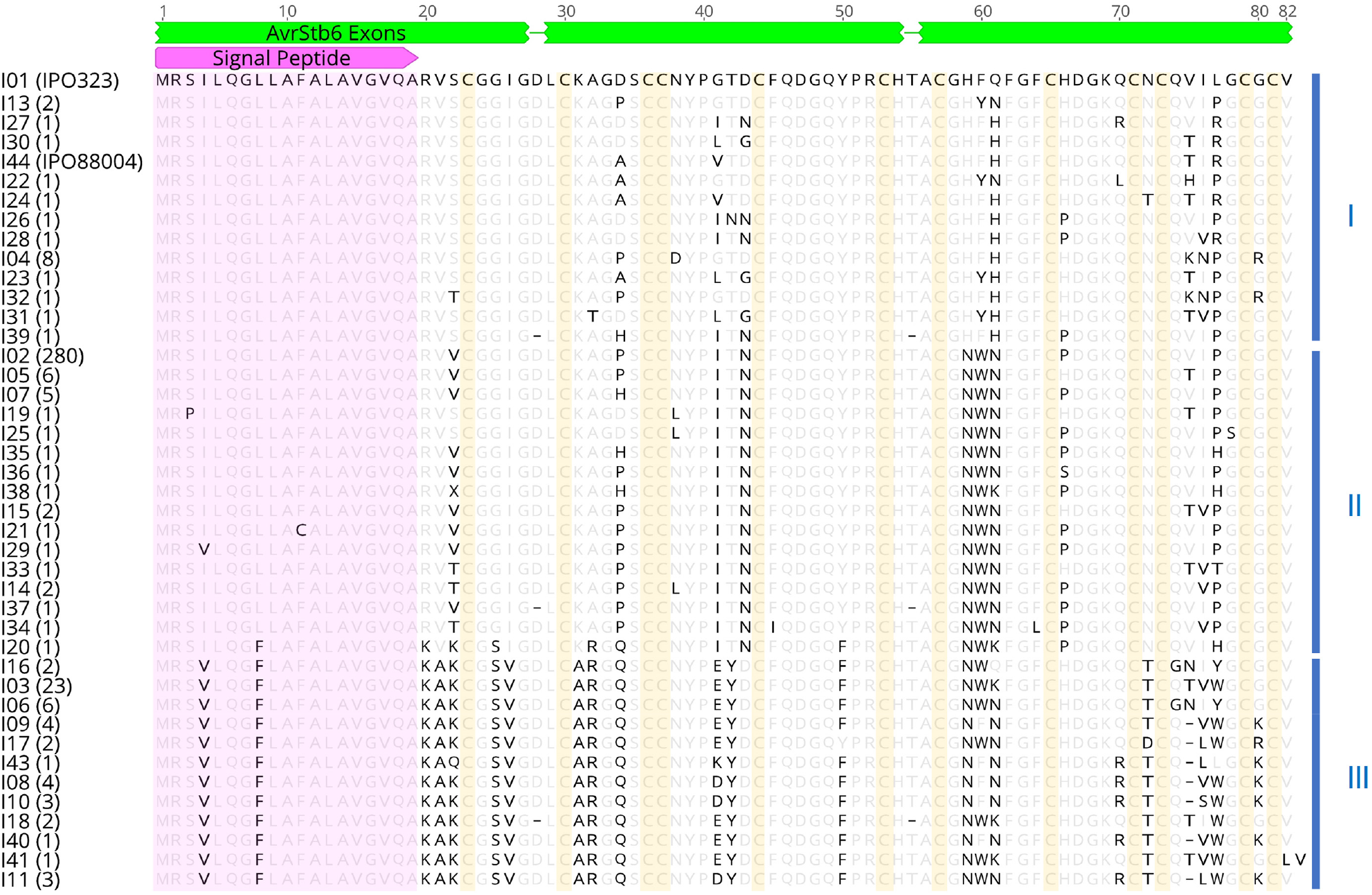
Alignment of the AvrStb6 isoform sequences identified in this study. Protein sequences were aligned using MAFFT v7.388. Numbers in brackets represent the number of isolates identified possessing each isoform. Isoforms I01 and I02 have been identified in the historic *Zymoseptoria tritici* isolates IPO323 and IPO88004 which are avirulent or virulent on *Stb6* wheat, respectively. Amino acids synonymous to the I01 sequence are greyed. Missing residues relative to I01 are represented as dashes. The pink arrow annotates the secretion signal peptide, whilst the green bars represent the exons of the coding gene sequence. The three alignment groupings (I, II and III) of the isoforms are indicated by vertical blue lines to the right of the sequences, and invariant cysteine residues are highlighted in yellow.

Almost all alleles were predicted to encode full-length AvrStb6 protein. The exceptions to this were A17 (only present in four isolates from Turkey) containing a nonsense mutation in the exon 1, and A24 and A46 (each represented by a single isolate also from Turkey) that contained single nucleotide deletions/frameshift mutations located in exon 3 and exon 2, respectively.

Twenty-nine alleles and twenty-five AvrStb6 protein isoforms each were uniquely present in single isolates in the collection. The majority of recovered *AvrStb6* sequences (361 of 381) were 365-bp in length from the ATG start codon to the TGA stop codon and including two introns, with others ranging in lengths from 362-bp to 366-bp. Most of the nucleotide polymorphisms were in exons 2 and 3, with exon 1 being more conserved, and particularly within the sequence coding for the N-terminal signal peptide (Figure 1). Eight alleles (A09, A10, A12, A21, A22, A23, A37 and A51) represented by 18 isolates contained three types of in-frame 3-bp deletions in exon 3. These alleles collapse to 7 protein isoforms (I08, I09, I10, I11, I17, I40 and I43). All these isoforms apart from I17, represented by two isolates from Western Europe, were exclusively found among the *Z. tritici* isolates from Turkey. The identified AvrStb6 isoforms comprised of 80–82 amino acid residues. From these, residues at 52 positions, including all 12 cysteines, were invariant across the collection (excluding the I42 frameshift mutant) (Figure 1). All other amino acid residues in AvrStb6 were variable, with the highest variation identified at positions 41, 61 and 77. As revealed by the amino acid sequence alignment and allele network analysis, AvrStb6 isoforms fell generally into three groups denoted from I, II and III (Figures 1, 2a, and S1). This, in principle, was supported by a phylogenetic analysis, which also identified that group III alleles, largely represented by isolates of Turkish origin, are phylogenetically more distant from those in groups I and II (Figure 2b).

**Figure 2.**
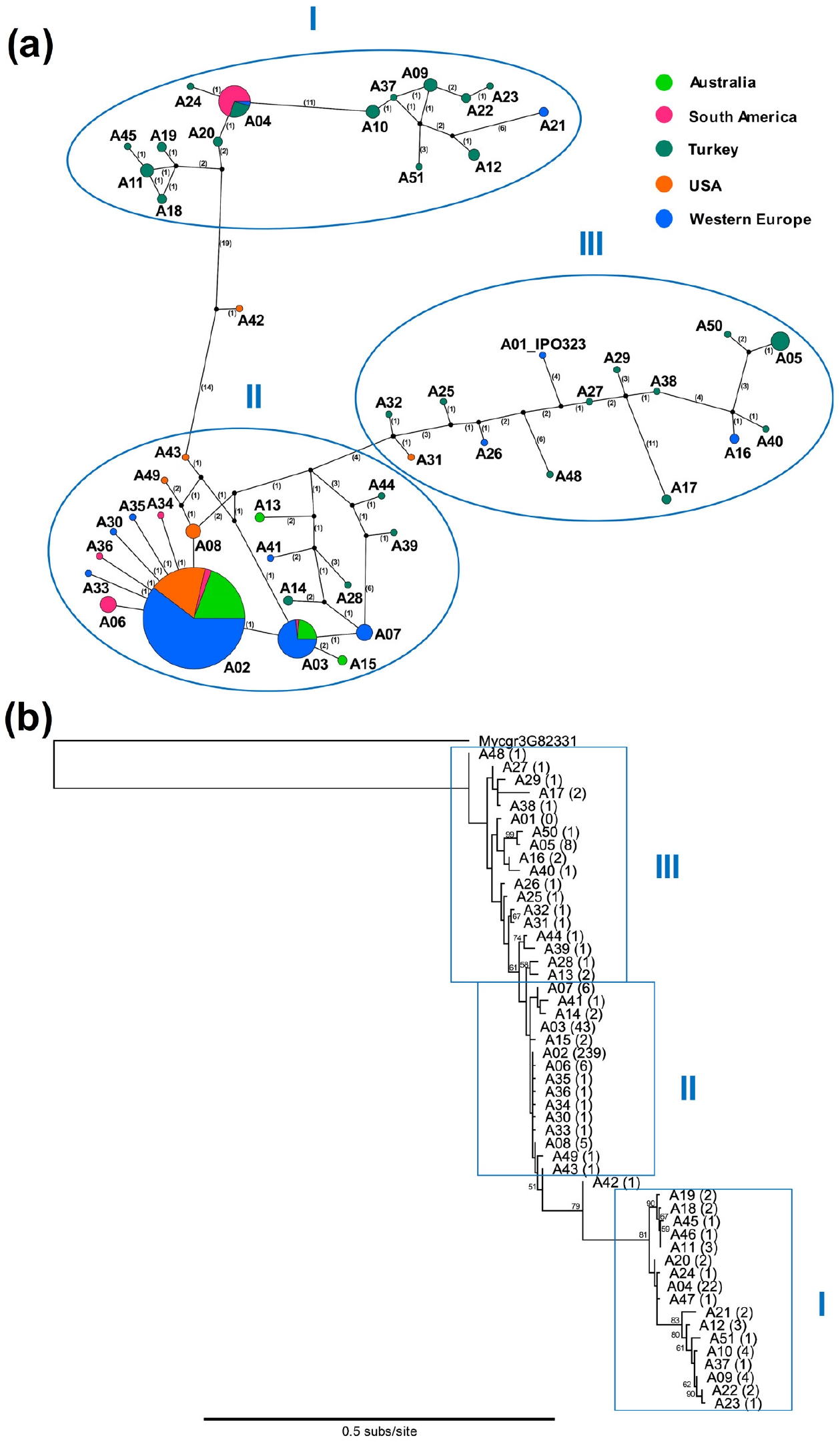
Analysis of the *AvrStb6* alleles distribution in global *Zymoseptoria tritici* population. (a) Allele network for global *AvrStb6* sequences obtained from the TCS analysis. Each node represents a separate allele identified in this study. The node size represents the number of isolates identified possessing the same allele. The nodes are painted in different colour depending on the geographic region(s) from which the isolates of the same node were identified. Connecting lines between nodes denote closely related alleles, with numbers in brackets corresponding to the number of mutations between adjoined alleles. Edges lengths are not proportional to genetic distances. Black unlabelled nodes represent hypothetical common ancestors of related alleles. The three groupings of alleles I, II and III are highlighted by blue ellipses. (b) Phylogenetic tree of the *AvrStb6* alleles constructed using PhyML. The tree was rooted using the *Z. tritici* gene *Mycgr3G82331* located on chromosome 10, a paralogue of *AvrStb6*. Numbers on branches indicate bootstrap scores for each branch, whilst numbers in brackets indicate the number of isolates identified possessing each allele.

### One AvrStb6 isoform predominates among current *Z. tritici* isolates globally

By far the most prevalent AvrStb6 isoform in the collection was I02, identified in 280 of 381 isolates (73.5%). This isoform was the most common almost in every geographic region studied including Australia (96.6%), Western Europe (91.8%), USA (82.7%) and Chile (80%) (Table 1). The only exceptions were Argentina and Uruguay where isoform I03 was most prevalent, and Turkey where isoform I02 was not identified at all. AvrStb6 isoform I02 was found to be encoded by several alleles: A02 (239 isolates), A03 (34 isolates), A06 (6 isolates), A15 (2 isolates), and A33 (1 isolate). However, in the USA part of the collection, only a single allele (A02) was found to code for I02.

**Table 1.** Prevalence of AvrStb6 isoform I02 in the recent global *Zymoseptoria tritici* collection *vs* in the historic collections.

| Region of Origin | Collection | Country/State of origin | Collection year(s) | Number of isolates | AvrStb6 I02 (%) |
| --- | --- | --- | --- | --- | --- |
| Western Europe | this study | England | 2015–2017 | 135 | 94.8 |
|  |  | Scotland | 2015–2017 | 14 | 92.9 |
|  |  | Ireland | 2015–2017 | 8 | 87.5 |
|  |  | France | 2015–2017 | 12 | 91.7 |
|  |  | Germany | 2015–2017 | 12 | 66.7 |
|  | Brunner and McDonald (2018) | Switzerland | 1999 | 29 | 51.7 |
|  | Zhong *et al.* (2017) | France | 2009–2010 | 102 | 49 |
| North America | this study | USA/Oregon | 2016 | 48 | 89.6 |
|  | Brunner and McDonald (2018) | USA/Oregon | 1990 | 56 | 0 |
| Australia | this study | Tasmania | 2014 | 58 | 96.6 |
|  | Brunner and McDonald (2018) | New South Wales | 2001 | 27 | 74.1 |
| South America | this study | Chile | 2016 | 10 | 90 |
|  |  | Argentina | 2016 | 10 | 20 |
|  |  | Uruguay | 2016 | 10 | 30 |
| Mediterranean | this study | Turkey | 2013–2016 | 46 | 0 |

The next most common isoform globally was I03 - the only other isoform to be found in multiple regions - which was identified in 23 isolates (6% of isolate collection) in Western Europe, South America and Turkey. All other identified AvrStb6 isoforms were represented by between 1 to 8 isolates in the collection, and remarkably well over half of the identified AvrStb6 isoforms (*n* = 25) were each recovered from single isolates (Figure 1).

### Turkey is a hotspot of *AvrStb6* diversity

Fifty-seven *Z. tritici* isolates were sourced from smallholder farmers’ wheat fields in Turkey as part of this study. From this sample, 29 *AvrStb6* alleles corresponding to 26 protein isoforms were identified. This represents a far higher rate of gene and protein sequence diversity than found in any other region studied here, with more than half of all identified AvrStb6 isoforms being found in Turkey and 25 being unique to this region. The network diagram of *AvrStb6* alleles (Figure 2a) visualises this high diversity, with both major branches largely consisting of alleles identified exclusively in the *Z. tritici* population from Turkey. Interestingly, isoform I02 - the most prevalent isoform globally - was not observed in the Turkish population. This population therefore represents a notable break in the trend described above.

### *Stb6* resistance-breaking isoforms are widespread in the current population

We next aimed to determine whether the various identified AvrStb6 isoforms conferred avirulence or virulence towards wheat genotypes carrying the cognate resistance gene *Stb6*. For this, for each of the nine AvrStb6 isoforms that were identified in more than one *Z. tritici* isolate in the collection (Figure S2) we randomly selected one isolate as representative for use in plant infection bioassays. One exception was the most common isoform I02, for which two representative isolates 2NIAB and R16.1 (collected in 2015 and 2016, respectively, from different commercial fields in the UK) were selected. The bioassays involved fungal inoculation of two pairs of wheat genotypes at the seedling stage, with the genotypes in each pair being genetically near identical but distinguished by the presence/absence of *Stb6*. One pair comprised wheat landrace Chinese Spring carrying the resistance-conferring haplotype 1 of *Stb6* (Saintenac *et al.*, 2018) and a near-isogenic line that contains the susceptibility haplotype 3 of *Stb6* developed from a cross between Chinese Spring and a susceptible cv. Courtot. The second pair comprised wheat cv. Cadenza (*Stb6*) and a mutant of cv. Cadenza that contains no functional *Stb6* due to a large CRISPR/Cas9-induced frameshift deletion in the first exon of this gene. The previously characterised avirulent and virulent *Z. tritici* isolates IPO323 (Brading *et al.*, 2002) and IPO88004 (Saintenac *et al.*, 2018) possessing isoforms I01 and I44, respectively, were used as controls. Of the ten isolates (representing 9 different isoforms) tested, nine induced typical STB disease symptoms and fungal pycnidia on all tested wheat genotypes irrespectively of the presence/absence of *Stb6* (Figures 3 and S3). This included the isolate(s) representing I02, the most prevalent AvrStb6 isoform in our collection.

**Figure 3.**
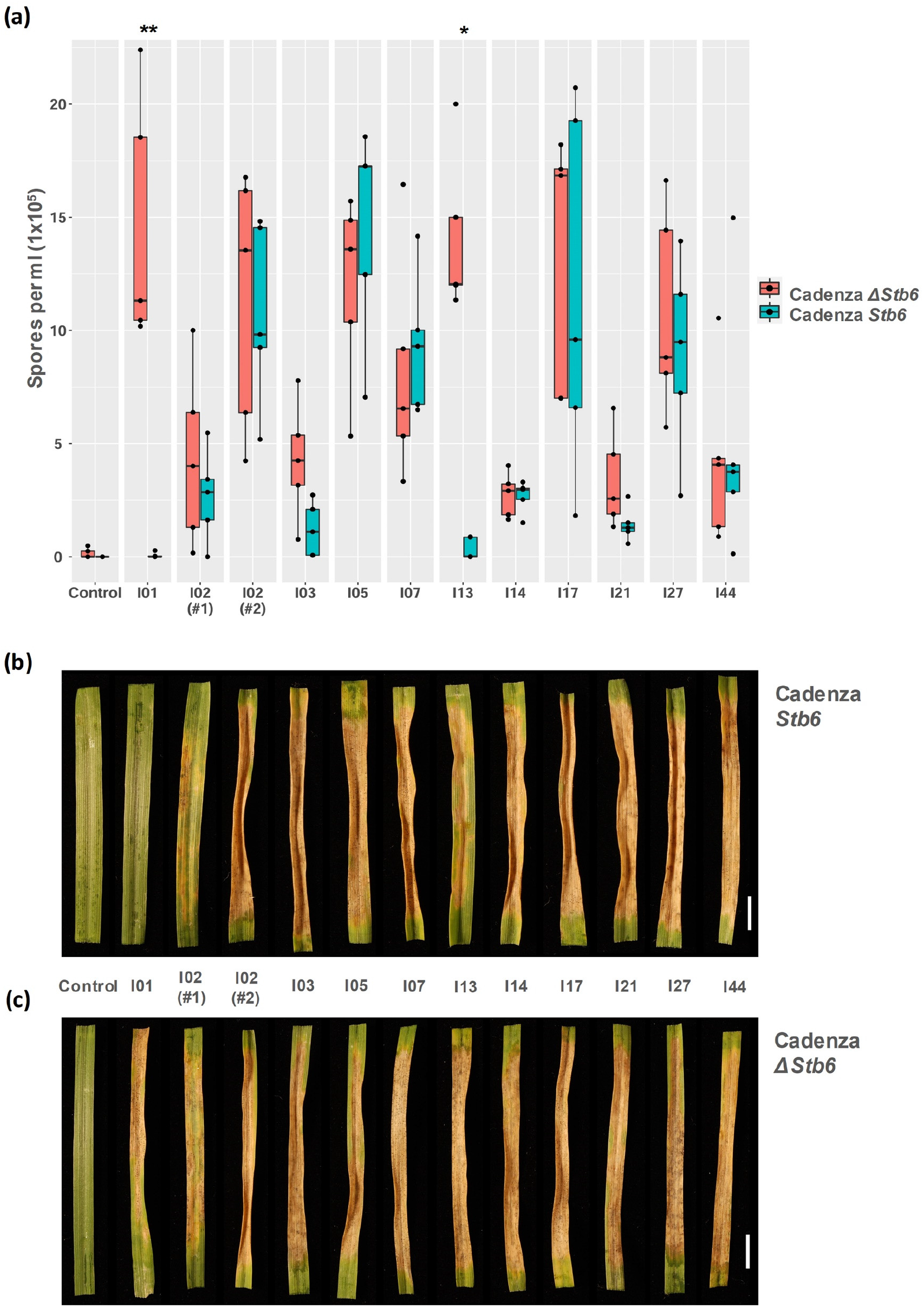
Plant inoculation bioassay. Leaves of differential wheat genotypes, wild-type cv. Cadenza containing *Stb6* and a CRISPR/Cas9-induced deletion mutant (Cadenza *ΔStb6*) that lacks the functional resistance gene in the same genetic background, were inoculated as young, three-week-old seedlings with a selection of *Z. tritici* isolates possessing different isoforms of AvrStb6. (a) Counts of pycnidiospores washed off the inoculated wheat leaves at 21 days post inoculation (dpi). Asterisks represent isolates with significant (*, *p* < 0.05) or highly significant (**, *p* <0.005) differences in pycnidiospore counts between the resistant and susceptible genotypes. The most common isoform I02 was represented by the two isolates (labelled #1 and #2) originating from two different geographic regions. (b) and (c) are images of inoculated wheat leaves harvested at 21 dpi and incubated for two days under ~ 100% humidity to induce pycnidiation. Scale bar, 10 mm.

Other than the avirulent reference isolate IPO323 (I01), only one other isolate could potentially be classified as avirulent on wheat possessing *Stb6,* as it induced development of some chlorosis but no necrosis nor any asexual sporulation structures (pycnidia) (Figure S3). This isolate possessed the AvrStb6 isoform I13, which shares the highest percent amino-acid identity with the known avirulence isoform I01 (Figure S2). Several isolates, possessing AvrStb6 isoforms classified as virulence, induced lower level of disease on the *Stb6*-containing wheat compared to their near isogenic genotypes that lack *Stb6* as reflected by differences in the observed pycnidospore counts (Figure 3). However, these differences were significant (at *p* < 0.01) only for the *Z. tritici* isolate T1-A2017.24 possessing the AvrStb6 isoform I13 and the control avirulent isolate IPO323.

### Avirulence isoforms of AvrStb6 are common on wheat genotypes irrespective of the *Stb6* resistance gene status

We wondered whether overrepresentation of AvrStb6 virulence isoforms in our isolates collection could have been due to most isolates potentially sampled from resistant, *Stb6-* containing wheat cultivars. The cultivar information was available for 254 out of 381 sampled *Z. tritici* isolates, and it was possible to obtain 20 cultivars - sources of 210 isolates - for further study (Table S1). Ten of these cultivars have been characterized in the previous study (Saintenac *et al.*, 2018) and their *Stb6* haplotype status and reaction to the avirulent strain IPO323 of *Z. tritici* are known (Table S1). We then re-sequenced *Stb6* following the procedure described in Saintenac *et al.* (2018) from the other ten cultivars including Consort, Dickens, Gallant, Genesis INIA 2375, Kaseberg, KWS Cashel, KWS Lumos, Marston, Solace, and Zulu to identify their *Stb6* haplotype status (Table S1). It was determined that 9/20 of these cultivars possess disease resistance conferring haplotype 1 of *Stb6* and they were sources of 73 (34.8 %) Z*. tritici* isolates in our collection, whereas the other 11/20 analysed cultivars contained non-functional susceptibility haplotypes 3 or 7 of *Stb6* and they represented sources of 137 (65.2 %) isolates (Table 2). Therefore, it appears that there was no overall bias in the wheat cultivars used in this study towards susceptible or resistant, and that a larger proportion of *Z. tritici* isolates was sampled from cultivars containing no functional *Stb6*.

**Table 2.**
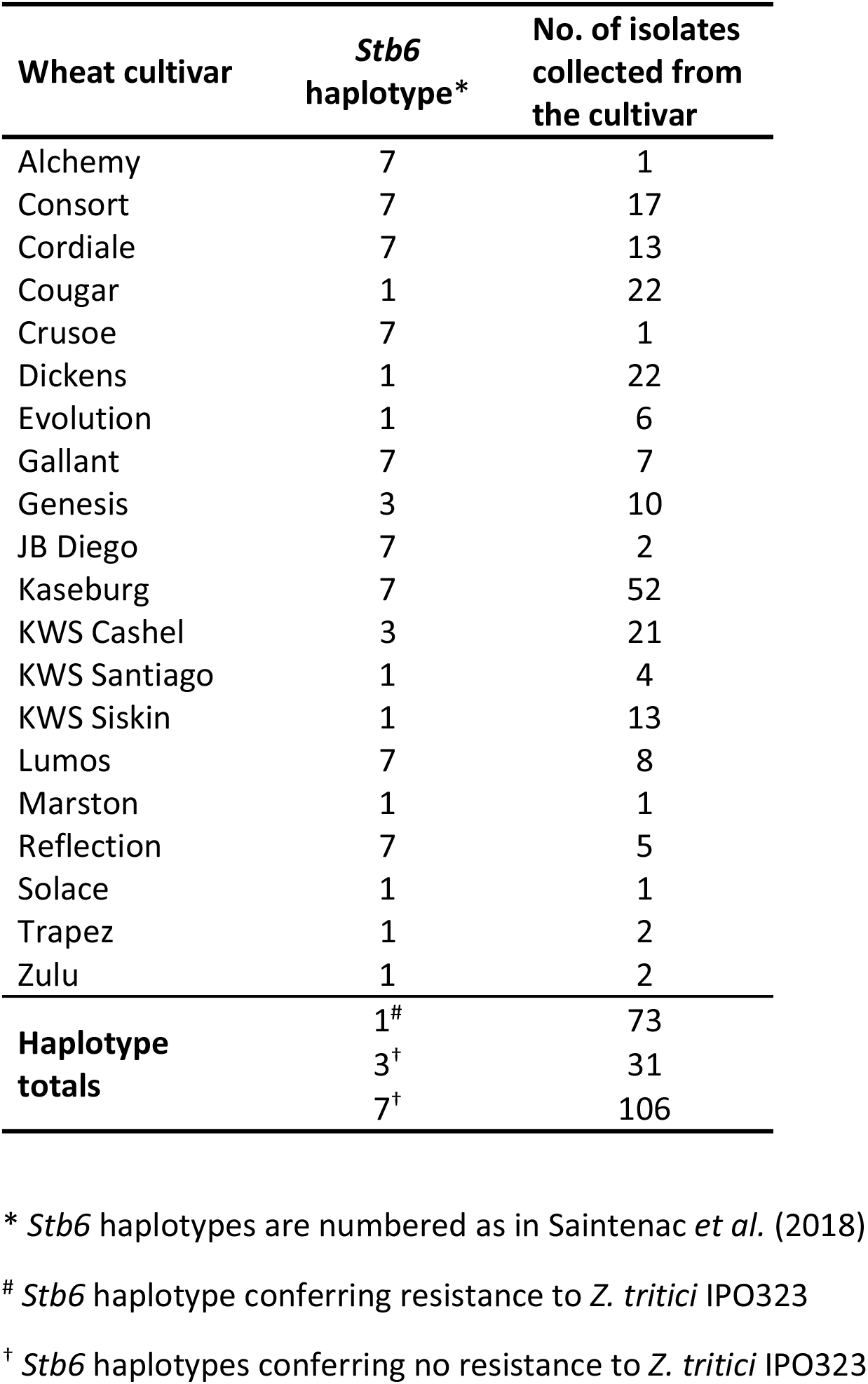
*Stb6* haplotype data for wheat cultivars used in this study.

### *AvrStb6* expression is highly variable between *Z. tritici* isolates

Our main hypothesis is that the breakdown of *Stb6*-mediated resistance by the current *Z. tritici* isolates analysed in this study (Figures 3 and S3) was due to the numerous changes in the AvrStb6 protein sequence identified (Figure 1). However, we also considered an alternative possibility whereby breakdown of *Stb6* resistance may have been due to the suppression of *AvrStb6* expression during infection. We therefore used a RT-qPCR to compare the expression levels of *AvrStb6* in the ten selected *Z. tritici* isolates (representing ten different most frequent AvrStb6 isoforms) that were virulent on *Stb6* wheat *versus* an avirulent isolate IPO323 during infection of a fully susceptible wheat cv. Taichung 29 that does not possess *Stb6* (Ghaffary *et al.*, 2012). As each of the 11 tested isolates has different infection dynamics on this wheat cultivar, the leaf tissues for use in RT-qPCR were sampled at different time points post fungal inoculation, depending on the isolate, when the first symptoms of the disease become visible. This infection phase was chosen as many *Z. tritici* effector genes including *AvrStb6* have previously been shown to display maximal expression during this transition to necrotrophy phase (Rudd *et al.*, 2015; Zhong *et al.*, 2017). RT-qPCR analysis revealed that *AvrStb6* was expressed in each tested isolate (Figure S4). Moreover, although the *AvrStb6* expression levels were highly variable between different isolates, in none of the virulent isolates with the possible exception of WAI1822 carrying AvrStb6 isoform 14 they were substantially lower than in an avirulent isolate IPO323 (Figure S4). This argues against the mechanism of *Stb6* resistance breakdown being due to suppression of *AvrStb6* expression.

## DISCUSSION

Few studies have analysed the temporal changes in Avr factor prevalence driven by selection pressure from host species in pathogen populations. Often, these have a fairly narrow geographical area coverage. For example, studies into the changes in *Avr* gene prevalence in the fungal pathogen *Leptosphaeria maculans* of *Brassica* crops over time have focussed on single countries such as Canada (Fernando *et al.*, 2018) or Australia (Van de Wouw *et al.*, 2018). On the other hand, studies with a global geographical scope often do not consider the impact of sampling time on changes in Avr factor prevalence. Here we analysed the diversity of the avirulence factor *AvrStb6* in the recent global *Z. tritici* populations. Moreover, we took advantage of the two published similar studies which analysed the population diversity of *AvrStb6* in the more historic isolate collections sampled in 1990–2001 or 2009–2010 (Zhong *et al.*, 2017; Brunner and McDonald, 2018). Intercomparing our data that was based on the analysis of isolates sampled in 2013–2017 with that from these two previous studies revealed that large shifts in *AvrStb6* allele prevalence have taken place in multiple global regions over a relatively short time period. These include the avirulence isoform of AvrStb6 originally described from the reference isolate IPO323 (collected in the Netherlands in 1981) becoming potentially extinct in the recent *Z. tritici* populations, and emergence of multiple virulence isoforms with one becoming predominant globally. We propose that these changes in *Z. tritici* populations are imposed by the worldwide extensive deployment of commercial wheat cultivars carrying *Stb6* in recent years (Chartrain *et al.*, 2005; Saintenac *et al.*, 2018).

The high proportion of AvrStb6 isoforms to alleles illustrates the high frequency of non-synonymous mutations within the effector gene, which in turn suggests a strong selection pressure for the evolution of novel AvrStb6 protein isoforms that cannot be recognised by the cognate wheat immune receptor Stb6. This phenomenon is well documented for other pathogen effectors, for instance with diversifying selection observed for ToxA, Tox1 and Tox3 of *Parastagonospora nodorum* (Stukenbrock and McDonald, 2007; McDonald *et al.*, 2013), NIP1 of *Rhynchosporium commune* (Schürch *et al.*, 2004), and AvrP123 and AvrP4 of the flax rust pathogen *Melampsora lini* (Barrett *et al.*, 2009). In several fungal pathogens, repeat-induced point (RIP) mutations have been shown to be responsible for the evolution of virulence due to the diversification of effector gene sequences including the introduction of multiple premature stop codons (Fudal *et al.*, 2009). Given the transposon rich, highly dynamic region of the genome in which *AvrStb6* is located (Sánchez-Vallet *et al.*, 2018) it seemed possible that a similar mechanism has contributed to the diversification of this *Z. tritici* effector. However, an *in-silico* analysis suggests a low frequency of RIP-associated mutations in the *AvrStb6* genomic region (Figure S5). Moreover, we found that the secretion signal peptide region and all the cysteine residues in AvrStb6 were highly conserved (Figure 1), whilst premature stop codon mutations within the gene were extremely rare. The signal peptide of effectors is required for their secretion into the host apoplast, whilst the formation of disulphide bonds between cysteines is thought to help to stabilise the protein in the harsh alkaline pH apoplastic environment. In addition, using RT-qPCR analysis we showed that *AvrStb6* was expressed in a set of *Z. tritici* isolates representing the most frequently occurring isoforms of this effector during a phase transition from biotrophy to necrotrophy (Figure S4). Whilst efforts to demonstrate a role for AvrStb6 isoforms in virulence have to date proved unsuccessful (Kema *et al.*, 2018), these findings combined suggest an evolutionary pressure to maintain a functional effector, implying potential contribution of AvrStb6 to fungal virulence or fitness, perhaps under certain as yet unknown environmental conditions.

Our study revealed the dominance of a single AvrStb6 isoform, I02, in the global population of *Z. tritici*. This isoform was found to be predominant in all countries of Western Europe from which the isolates were sourced, as well as Australia, USA and Chile. It is unknown why this isoform specifically became so prevalent in multiple regions around the world recently, given the ability of isolates carrying other isoforms to also evade *Stb6*-mediated resistance (Figure 3). However, the limited diversity of *Stb6* haplotypes (*n* = 3) in the host wheat plants sampled (Table 2) may suggest that AvrStb6 isoform I02 is specifically adapted for successful infection of wheat possessing these *Stb6* haplotypes. The virulence isoform I02 may also be the best-adapted for carrying out its as yet unknown effector function. Notably, sequencing of *AvrStb6* from *Z. tritici* isolates sourced from Oregon, USA in 1990 failed to identify a single occurrence of I02 (Table 1). This is in sharp contrast to our study which identified prevalence of this isoform, found in 82.7% of the current isolates in the Oregon part of our collection. This population shift, combined with the fact that in all cases I02 was coded by a single allele A02, may suggest the introduction and rapid proliferation of this virulence *AvrStb6* allele throughout the population of *Z. tritici* in Oregon, as occurred for example with the recent accidental importation of *Pyricularia oryzae*, a wheat blast pathogen, from Brazil to Bangladesh (Islam *et al.*, 2016).

Our data also revealed a profound increase in the prevalence of AvrStb6 isoform I02 in the global recent (2013–2017) *Z. tritici* population compared to that in the older collections comprising isolates sampled in 1990–2001 (Brunner and McDonald, 2018) or 2009–2010 (Zhong *et al.*, 2017) (Table 1). One exception to this is Australia, where a prevalence of AvrStb6 I02 in the local *Z. tritici* population sampled in 2001 was already evident (Zhan *et al.*, 2005) (Table 1). Interestingly, in addition to the increasing dominance of a single isoform I02, the overall diversity of AvrStb6 isoforms seem to have decreased over time. Thus, for example, previous studies of historic isolates by Brunner and McDonald (2018) and Zhong *et al.* (2017) identified thirty and eighteen AvrStb6 isoforms in the populations comprising 142 or 103 isolates, respectively. These numbers are substantially higher than the 42 isoforms identified from 381 isolates analysed in the current study. Such a large shift in the *AvrStb6* population diversity that occurred over a relatively short period of time and the increased fixation of AvrStb6 I02 in the population could have been caused by the decreased genetic diversity of wheat due to breeding (Fu and Somers, 2009) and widespread global deployment of *Stb6-*containing cultivars by arable farmers (Chartrain *et al.*, 2005; Saintenac *et al.*, 2018). Other studies have shown changes in the Avr factor prevalence, such as an increased incidence of *AvrLm4-7* in *Leptosphaeria maculans* from 47.2% in 2005–2006 (Dilmaghani *et al.*, 2009) to 89.7% in 2010–2011 (Liban *et al.*, 2016) in Western Canada, demonstrating the ability of pathogens to adapt rapidly to changing conditions in host populations. However, to our knowledge this is the first time that a shift of such significance has been shown to occur across a global population of a plant pathogen.

The analysis of *AvrStb6* alleles did not identify I02 as the dominant isoform in Argentina and Uruguay, and this isoform was completely absent from the Turkish *Z. tritici* population. It is possible that the local wheat cultivars used for sampling *Z. tritici* isolates for this study in these countries may have unique genetic makeups, although it was not possible to test this hypothesis in this study. The Turkish population is of particular interest, given it has a substantially higher *AvrStb6* allele diversity compared to those in other wheat growing regions studied here. Whilst the exact reasons for this difference are unknown, we hypothesise that the widespread use of local genetically diverse landraces, instead of genetically more similar commercial wheat cultivars, in Turkey is a driver for matching diversification of *Z. tritici* genomes. The location of Turkey near or at the hypothesised geographic region of origin of wheat (Shewry, 2009) and *Z. tritici* (Stukenbrock *et al.*, 2006) may also be a cause for the high diversity of *AvrStb6* alleles observed here because origins of species often also correspond to the centres of their genetic diversity.

A recent study (Kema *et al.*, 2018) suggested that the *Z. tritici* pathogenicity towards wheat cultivars carrying the disease resistance gene *Stb6* was associated with changes at the amino acid positions 41 and 43 in the AvrStb6 protein. The new data obtained here support these findings and we confirm that isolates possessing isoforms I01 and I13 having G and D at positions 41 and 43, respectively (Figure S2), were unable to cause disease on wheat cultivars possessing *Stb6* (Figure 3). However, we also found that the *Z. tritici* isolates expressing AvrStb6 isoform I03, I17, or I44, having E or V instead of G at position 41 but D at position 43 were fully virulent on *Stb6* wheat (Figure S2). This suggests that amino acid changes in AvrStb6 at position 41 alone may be responsible for the *Stb6* resistance breaking ability of *Z. tritici* isolates. It would be interesting to test this hypothesis through targeted mutagenesis in follow on studies.

The number of *Z. tritici* isolates analysed in this study and the failure to identify AvrStb6 isoform I01 in the collection, suggest that this isoform has been eliminated from the modern *Z. tritici* population worldwide. However, the *AvrStb6* allele network analysis indicates that the true diversity of *AvrStb6* may be higher than identified as in some cases multiple mutations were found to separate the neighbouring alleles in the network (Figure 2a). A significant amount of undiscovered sequence variation in *AvrStb6* may still be present in the global *Z. tritici* population. However, one of the main conclusions from our study is that the wheat disease resistance gene *Stb6* has currently been almost completely overcome due to predominance of fungal isolates carrying resistance breaking AvrStb6 isoforms in the modern *Z. tritici* populations around the world. This suggests, that despite the ability of *Z. tritici* to sexually reproduce on resistant hosts (Kema *et al.,* 2018), Avr factors tend to be eliminated in subsequent populations.

## EXPERIMENTAL PROCEDURES

### *Zymoseptoria tritici* isolates collection

The 381 *Z. tritici* isolates used in this study were collected from naturally infected wheat fields in the UK (*n* = 150), Ireland (*n* = 8), France (*n* = 12), Germany (*n* = 12), Australia (*n* = 58) and Turkey (*n* = 57), and from single fields in Chile (*n* = 10), Argentina (*n* = 10), Uruguay (*n* = 10) and USA (*n* = 52). Most of these isolates were collected between 2015–2017, with the exception of isolates collected in Australia (2014) and Turkey (2013–2016), and the control isolates IPO323 (avirulent on *Stb6* wheat) and IPO88004 (virulent on *Stb6* wheat) collected in 1981 and 1988 in the Netherlands and Ethiopia respectively (Kema and van Silfhout, 1997; Goodwin *et al.*, 2011). Stocks of *Z. tritici* isolates were stored as blastospore water suspensions in 50% (v/v) glycerol at −80°C.

### Sequence, diversity, and phylogenetic analysis of *AvrStb6*

Genomic DNA extraction was carried out from *Z. tritici* isolates as previously described (Rudd *et al.*, 2010). Primers for PCR amplification and sequencing of the *AvrStb6* gene were designed in the upstream and downstream UTRs of the gene using publicly available and own whole-genome sequencing data for European *Z. tritici* isolates (Chen *et al.,* Rothamsted Research, Harpenden, UK, 2019, personal communication). Primers avrstb6.f1 and avrstb6.r1 were selected for amplification of *AvrStb6* from European isolates, and primers avrstb6.f3 and avrstb6.r1 were used to amplify *AvrStb6* from North and South American isolates (Table S2). All amplifications were done using Phusion High-Fidelity DNA Polymerase (New England Biolabs UK Ltd., Hitchin, UK). PCR products were purified using a QIAquick PCR Purification Kit (Qiagen UK Ltd., Manchester, UK) and Sanger sequenced in house or at Eurofins Genomics UK Ltd., Wolverhampton, UK. *AvrStb6* sequences from the Australian *Z. tritici* isolates were extracted from the whole-genome sequencing data (NCBI BioProject accession number PRJNA480739; McDonald *et al.*, 2019).

*AvrStb6* allele and the corresponding protein isoform sequences were aligned using MAFFT v.7.388 (Multiple Alignment using Fast Fourier Transform; Katoh and Standley, 2013), and the phylogenetic tree was generated from the allele alignment using PhyML v.3.3.20180621 (Phylogenetic inferences using Maximum Likelihood; Guindon and Gascuel, 2003) with 1000 bootsraps. The *AvrStb6* sequence from the *Z. tritici* isolate IPO323 was used as a reference for MAFFT alignments, whilst a paralogue of *AvrStb6* present on chromosome 10 (gene *Mycgr3G82331*; Brunner and McDonald, 2018) was used to root the phylogenetic tree. A TCS allele network (Clement *et al.*, 2000) was created to visualise the diversity of *AvrStb6* alleles identified in the population using PopArt 1. 7 (Leigh and Bryant, 2015). Coding regions were annotated and translated to amino acid sequence using Geneious 10.2.3 (Biomatters Ltd., Auckland, New Zealand). Repeat-induced point mutation (RIP) analysis of the region of *Z. tritici* chromosome 5 containing *AvrStb6* was carried out using RIPper software (van Wyk *et al.,* 2019).

The *AvrStb6* sequence diversity data obtained here was compared to the similar previously published datasets for the historic *Z. tritici* collections. The *AvrStb6* allele sequences obtained in Zhong *et al.* (2017) were kindly provided by Daniel Croll (University of Neuchâtel, Switzerland) and those obtained in Brunner and McDonald (2018) were downloaded from the NCBI PopSet Database (accession number 1337388362).

### Generation of wheat genetic material

A wheat Chinese Spring near-isogenic line that carries the susceptibility haplotype 3 at the *Stb6* locus was obtained following five backcrosses starting with F_1_ Chinese Spring × Courtot. In each generation, plants were assessed for their responses to the avirulent *Z. tritici* isolate IPO323 and genotyped with simple sequence repeat marker GWM369 to maintain the susceptibility haplotype of *Stb6* in the progenies. A single BC_5_F_1_ plant heterozygous at the *Stb6* locus was self-pollinated. BC_5_F_2_ plants either homozygous for the susceptibility haplotype of *Stb6* were selfed and constitute the near-isogenic line used in this study.

The Cadenza *ΔStb6* mutant was produced by targeting the first exon of the *Stb6* gene with CRISPR/Cas9 at five sites (Figure S6). The sgRNA constructs, carrying two sgRNAs each (annotated sequences are available on figshare), were assembled using the pENTR4-sgRNA4 vector as previously described (Zhou *et al.*, 2014). All three sgRNA plasmids were co-delivered along with pCas9-GFP (Zhang *et al.*, 2019) encoding the wheat codon-optimised Cas9 fused in frame at the C-terminus with Green Fluorescent Protein (GFP), and pRRes1.111 (Alotaibi *et al.*, 2018) which directs expression of the *bar* selectable marker gene into immature wheat embryos of wheat cv. Cadenza using biolistics essentially as described in Sparks and Doherty (2020). CRISPR/Cas9 mutagenized T_0_ lines were genotyped for indels using the PCR band-shift assay (Nekrasov *et al.*, 2017). Cas9, sgRNA and *bar* transgenes were segregated out in subsequent generations and the homozygous Cadenza *ΔStb6* T_2_ line was used for phenotyping with *Z. tritici*.

### Fungal bioassays

Leaves of three-week-old wheat seedlings were inoculated with *Z. tritici* blastospore suspensions at 1 x 10^7^ spores/mL as previously described (Keon *et al.*, 2007). Wheat landrace Chinese Spring and cv. Cadenza, both carrying *Stb6* were used as resistant controls. A near-isogenic line in the Chinese Spring background that carries a susceptibility haplotype at *Stb6* locus, and a CRISPR/Cas9-induced Cadenza *ΔStb6* mutant were used as susceptible controls. A split plot randomised experimental design was used when carrying out the bioassays.

STB disease symptoms on each inoculated leaf were visually assessed at 21 days post inoculation (dpi) as previously described (Lee *et al.*, 2015). To complement the visual disease assessment, we carried out pycnidiospore wash counts as follows. Approximately 6-cm-long segments were cut from the inoculated leaves and these were exposed to high relative humidity at 15°C for 48 hours in the dark to induce pycnidiation. Pycnidiospores were washed off the infected leaves by addition of 2 mL of distilled water supplemented with 0.01% Tween-20 followed by vortexing for 30 seconds. Optical density at 600 nm (OD_600_) of the harvested pycnidiospore suspensions were measured by spectrophotometer. This was converted to spores/mL by comparing against a standard curve produced by measuring OD_600_ of control suspensions containing defined pycnidiospore numbers as counted by using a haemocytometer.

### Wheat *Stb6* haplotype analysis

*Stb6* haplotype data for ten wheat cultivars, sources of the *Z. tritici* isolates used here, was available from the recently published study (Saintenac *et al.*, 2018). For ten other cultivars, with previously unknown *Stb6* status, we extracted genomic DNA from 3-week-old wheat plants and re-sequenced *Stb6* exons following previously published methodology (Saintenac *et al.*, 2018). For *Z. tritici* isolates from Australia, Turkey and a few from Western Europe, it was not possible to establish from which cultivar they were sampled.

### Analysis of *AvrStb6* expression during the infection of wheat

Three-week-old seedlings of wheat cv. Taichung 29, which is highly susceptible to *Z. tritici* and contains no known genes/QTLs for resistance to STB (Ghaffary *et al.*, 2012), were inoculated with *Z. tritici* spore suspension at 1 × 10^7^ spores/mL as previously described (Keon *et al.*, 2007). At the emergence of visible symptoms on the inoculated leaves between 12-16-dpi (depending on the particular isolate), the leaves were harvested and snap-frozen in liquid nitrogen. Total RNA was extracted using TRIzol (Fisher Scientific - UK Ltd, Loughborough, UK) following the manufacturer’s instructions, and any potential traces of genomic DNA were removed using TURBO DNA-free kit (Fisher Scientific - UK Ltd). First strand cDNA was synthesized from 1 µg of total RNA using SuperScript IV Reverse Transcriptase (Fisher Scientific - UK Ltd). First strand cDNA preparations were diluted 1:10 with RNase-free water and subjected to RT-qPCR using SYBR Green JumpStart Taq ReadyMix (Merck Life Science UK Ltd., Gillingham, UK), with primers specified in Table S2. Three biological and three technical replicates were carried out for each sample, with relative expression of *AvrStb6* to the *Z. tritici* housekeeping gene *glucose-6-phosphate 1-dehydrogenase* (*G6PDH*; gene ID Mycgr3G100879) calculated using the 2^-ΔΔCT^ method (Livak and Schmittgen, 2001).

## ACKNOWLEDGEMENTS

We are grateful to Suzanne Clarke for her help with the experimental design of plant inoculation bioassays; Maider Abadie and Ana Machado-Wood for their advice and assistance with the *AvrStb6* gene expression analysis; and Hongxin Chen for sharing his spectrophotometry-based method of estimating concentration of pycnidiospores in leaf-wash suspensions. Our gratitude goes to many other colleagues for providing materials used in this study, namely Guilherme Rossato-Augusti, and Pilar Diez for the genomic DNA preparations of the UK, European, USA and South American *Z. tritici* isolates; Daniel Croll for the *AvrStb6* alleles data for the historic *Z. tritici* population analysed in Zhong *et al.* (2017); Hongxin Chen for the unpublished whole-genome sequences of various European *Z. tritici* isolates; Lucy Hyde and Caroline Sparks for advice and help with wheat transformation; Keith Edwards for sharing the pCas9-GFP construct; Florence Cambon for her help with the development of a nearly-isogenic Chinese Spring wheat line carrying a susceptibility haplotype at the *Stb6* locus; Mike Hammond-Kosack for genomic DNA preparations of several wheat cultivars used for *Stb6* exons re-sequencing; Martin Quinke and Christina Hagerty for seeds of USA and South American wheat cultivars, respectively; and Javier Palma-Guerrero for the *Z. tritici G6PDH*-specific RT-qPCR primer sequences. This work was supported by the UK Biotechnology and Biological Sciences Research Council (BBSRC) Nottingham-Rothamsted Doctoral Training Programme grant (BB/M008770/1) and the Institute Strategic Program grant ‘Designing Future Wheat’ (BB/P016855/1).

## DATA AVAILABILITY STATEMENT

The sequencing data that support the findings will be openly available in GenBank at https://www.ncbi.nlm.nih.gov/nucleotide/ following an embargo from the date of publication, accession numbers: MT856831–MT856877. Annotated sgRNA construct sequences in the GenBank format are available on figshare at https://doi.org/10.6084/m9.figshare.12964589.v1.

## SUPPORTING INFORMATION

**Table S1.** *Zymoseptoria tritici* isolates used in this study.

**Table S2.** Primers used in this study.

**Figure S1.** Frequency of each of the identified *AvrStb6* alleles along with their geographic origin.

**Figure S2.** Alignment of AvrStb6 isoforms from *Zymoseptoria tritici* isolates tested in pathoassays to determine virulence on *Stb6* containing wheat.

**Figure S3.** Plant inoculation bioassay.

**Figure S4.** Expression levels of different *AvrStb6* alleles during *Zymoseptoria tritici* infection of wheat.

**Figure S5.** Analysis of repeat induced point mutation frequency at the *AvrStb6* locus on *Zymoseptoria tritici* chromosome 5 using RIPper (http://theripper.hawk.rocks).

**Figure S6.** Alignment of the part of the *Stb6* coding sequence (exon 1) with the corresponding region in the CRISPR/Cas9-induced wheat Cadenza *ΔStb6* mutant.

