## Supporting Information for "Remarkable recent changes in genetic diversity of the avirulence gene *AvrStb6* in global populations of the wheat pathogen *Zymoseptoria tritici*"

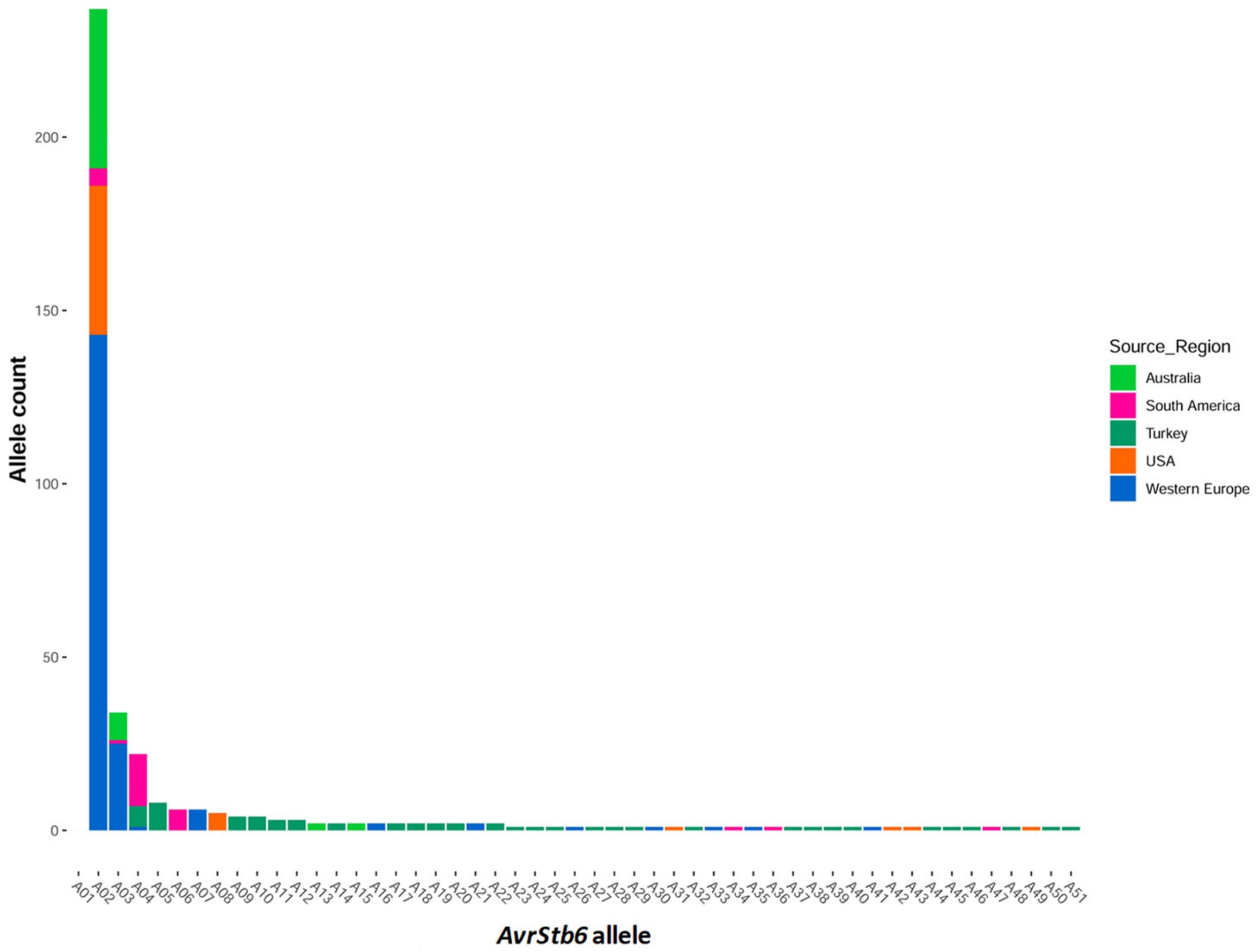


**Figure S1. Frequency of each of the identified *AvrStb6* alleles along with their geographic origin.**

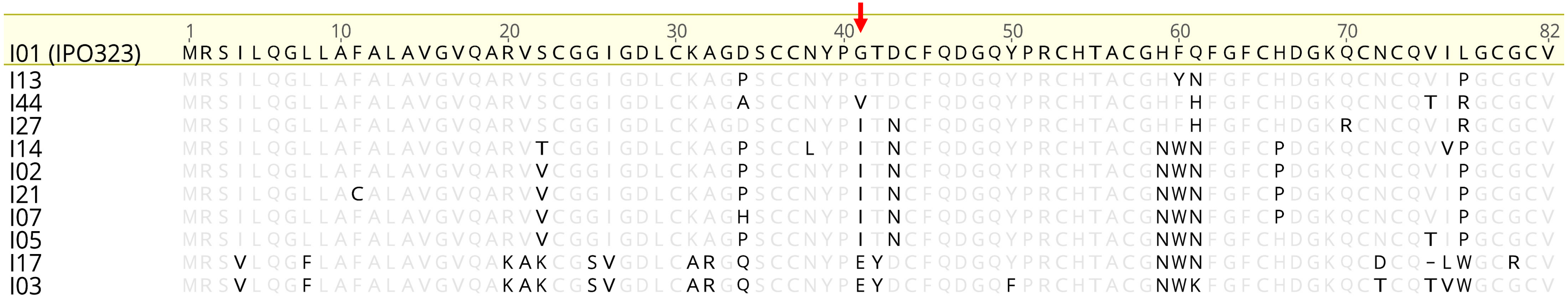


**Figure S2. Alignment of AvrStb6 isoforms from *Z. tritici* isolates tested in pathoassays to determine virulence on *Stb6* containing wheat**. Sequences were aligned using MAFFT v7.388. Of the above isoforms, all isolates were found to be virulent on *Stb6* containing wheat, with the exception of isolates possessing I01 (from reference isolate IPO323) and I13. Amino acids synonymous to the I01 reference sequence from isolate IPO323 are greyed. Missing residues relative to the reference are represented as dashes. The red arrow indicates the amino acid position #41, variation at which is hypothesised to determine the virulence/avirulence phenotype.

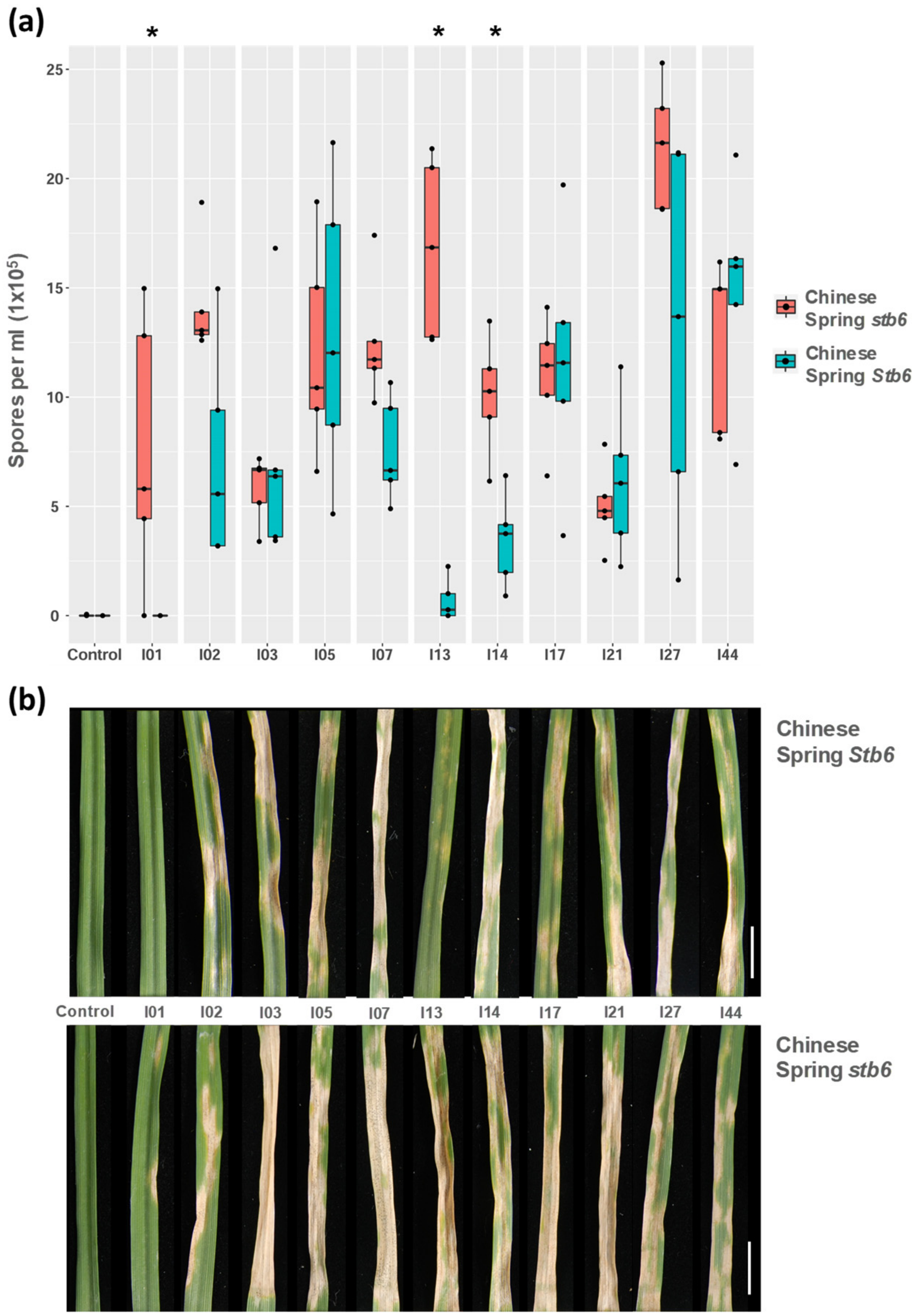


**Figure S3. Plant inoculation bioassay**. Leaves of differential wheat genotypes, landrace Chinese Spring (*Stb6*) and a nearly isogenic line developed from a cross with cv. Courtot carrying a susceptibility allele of *Stb6*, were inoculated as young, three-week-old seedlings with a selection of *Z. tritici* isolates possessing different isoforms of AvrStb6. (a) Counts of pycnidiospores washed off the inoculated wheat leaves at 21 days post inoculation (dpi). Asterisks represent isolates with significant (*, $p < 0.05$) or highly significant (**, $p < 0.005$) differences in pycnidiospore counts between the resistant and susceptible genotypes. (b) Images of inoculated wheat leaves harvested at 21 dpi and incubated for two days under ~ 100% humidity to induce pycnidiation. Scale bar, 10 mm.

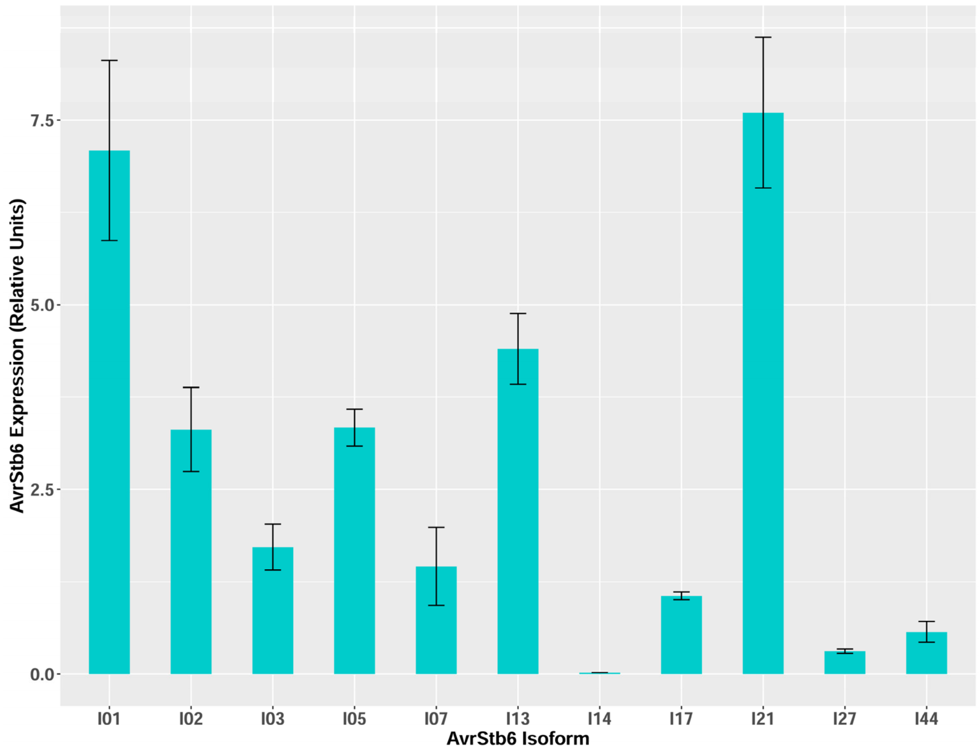


**Figure S4. Expression levels of different *AvrStb6* alleles during *Z. tritici* infection of wheat.** Leaves of highly susceptible wheat cv. Taichung 29, containing no known Septoria tritici blotch resistance genes, were inoculated with a selection of *Z. tritici* strains representing different *AvrStb6* alleles (giving rise to different protein isoforms) were harvested upon emergence of visible disease symptoms. Error bars are standard errors from three biological replicates. Expression levels are relative to the expression of the housekeeping gene *G6PDH* that encodes glucose-6-phosphate 1-dehydrogenase.

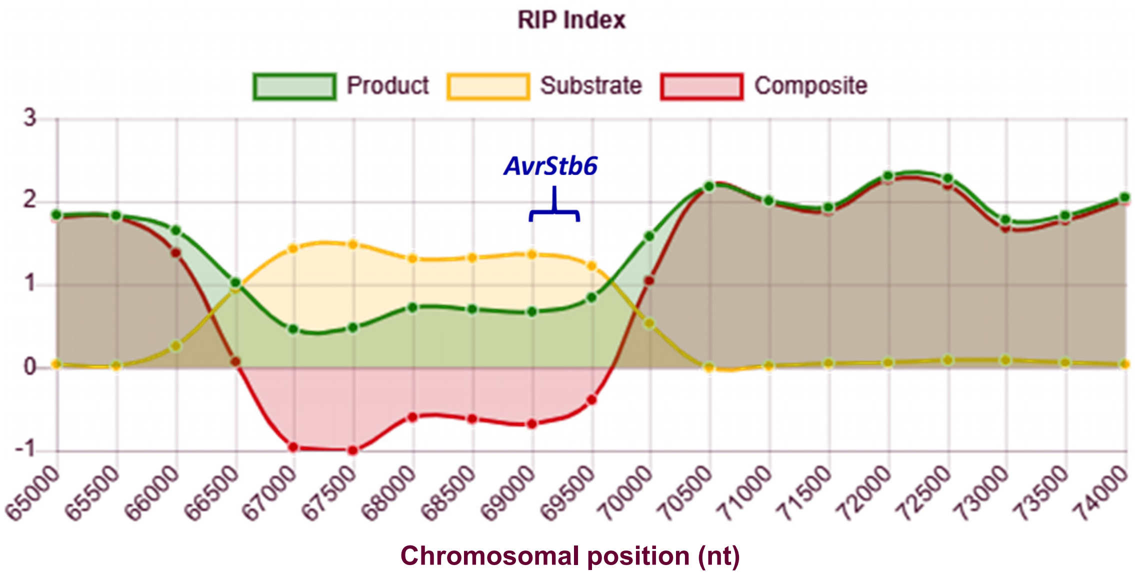


**Figure S5. Analysis of repeat induced point mutation frequency at the *AvrStb6* locus on *Zymoseptoria tritici* chromosome 5 using RIPper** (http://theripper.hawk.rocks). The *AvrStb6 gene* is located between nucleotides 69019 and 69383. 'Substrate' and 'Product' represent the frequency of RIP targeted dinucleotides (G and C) and the RIP product dinucleotides (T and A), respectively. 'Composite' represents the observed decrease in Substrate against the increase in Product, and provides an estimate of RIP frequency. See for software. Window size = 1000 bp, slide size = 500 bp.

```
                                                                target 48f
STB6 WT 1   ATGTCTCTGAGCTGCTGGTCCTGGTTCCTCGCCTTCGCCTGGGTTTGGTGTCTGCCACTG   60
 ΔStb6 1    ATGTCTCTGAGCTGCTGGTCCTGGTTCCTCGCCTTCGCCTGGGTTTGGTGTCTGCCACTG   60

                                                              target 104r
STB6 WT 61  ATGCTCATGGCGGCCGAGGAGCAGCAAGGGGATGGCTGCTTGGCCAAGTGTGGCAGCGTC  120
 ΔStb6 61   A------------------------------------------------GTGGCAGCGTC   72

STB6 WT 121 ACCATCTCCCCCCCGTTCTGGCTCACTGATTGGCAAACAGGAAGATTATGTGGTTCGCCT 180
 ΔStb6 73   ACCATCTCCCCCCCGTTCTGGCTCACTGATTGGCAAACAGGAAGATTATGTGGTTCGCCT 132

STB6 WT 181 GGACCGCTGGACTTCGAGCTTACATGCTATAACGGCAGTTATCCACTTCTTCCAAGCTCT 240
 ΔStb6 133  GGACCGCTGGACTTCGAGCTTACATGCTATAACGGCAGTTATCCACTTCTTCCAAGCTCT 192

                                                     target 278r
STB6 WT 241 GTGCCCAACAACGCCGGCTTTGCAATCATGGACATATCCTATGAGGAACGCAGCTTGCGC 300
 ΔStb6 193  GTGCCCAACAACGCCGGCTTTGCAATCATGGACATATCCTATGAGGAACGCAGCTTGCGC 252

STB6 WT 301 GTCGTTGATCTACGCAAGCTGCAACTATTACACGACCCGCCCAACATCTTCAACAGCTGC 360
 ΔStb6 253  GTCGTTGATCTACGCAAGCTGCAACTATTACACGACCCGCCCAACATCTTCAACAGCTGC 312

STB6 WT 361 TTGCCGATGTGGAACACCTCTGCCAAACTGGGCCGCCCGTTTAAGATCTCCCCCGTCAAC 420
 ΔStb6 313  TTGCCGATGTGGAACACCTCTGCCAAACTGGGCCGCCCGTTTAAGATCTCCCCCGTCAAC 372

                                                         target 465f
STB6 WT 421 CTGGAACTCATCTTGTACAACTGCACGGAGAAGGCCGCCGCGGCGGCACGCCTGGATAAA 480
 ΔStb6 373  CTGGAACTCATCTTGTACAACTGCACGGAGAAGGCCGCCGCGGCGGCACGCCTGGATAAA 432

STB6 WT 481 GAACTGGTGCAGGCGAAGACGATGAGGTGCGTGAACACAAGCAACACGTTTGTTCATGCG 540
 ΔStb6 433  -AACTGGTGCAGGCGAAGACGATGAGGTGCGTGAACACAAGCAACACGTTTGTTCATGCG 491

STB6 WT 541 GGGGTGCCATACGACACCACCGGGACCTACTCTAGTTATGCTTTGGAGGGCTGCGTTCCA 600
 ΔStb6 492  GGGGTGCCATACGACACCACCGGGACCTACTCTAGTTATGCTTTGGAGGGCTGCGTTCCA 551

STB6 WT 601 ATCGTCTTGCCGGTGCTGCGCTTGCCATCCGGCGAGACGAACACGAGCCACTACGAGCGG 660
 ΔStb6 552  ATCGTCTTGCCGGTGCTGCGCTTGCCATCCGGCGAGACGAACACGAGCCACTACGAGCGG 611

STB6 WT 661 CTCATCCAAAGTGGCTTCCTCCTGAAATGGGAACTGCCCCCTCCTCTCCCTGCACCTGCA 720
 ΔStb6 612  CTCATCCAAAGTGGCTTCCTCCTGAAATGGGAACTGCCCCCTCCTCTCCCTGCACCTGCA 671
              target 715r
STB6 WT 721 CCTAGGAATGAACCACCTCCCCCTCCAG 748
 ΔStb6 672  CCTAGGAATGAACCACCTCCCCCTCCAG 699
```

**Figure S6. Alignment of the part of the *Stb6* coding sequence (exon 1) with the corresponding region in the CRISPR/Cas9-induced wheat Cadenza *ΔStb6* mutant.** Induced deletions are shaded in red. sgRNA targets are shown in blue. Protospacer adjacent motifs (PAMs) are shown in green. The premature STOP codon in *ΔStb6* is shown in red.

**Table S1.** *Zymoseptoria tritici* isolates used in this study.

| Isolate code | Isolate name | Region of origin | Country of origin | Year collected | Stock held | Wheat cultivar source | Wheat cultivar *Stb6* haplotype | *AvrStb6* allele[#] | AvrStb6 isoform | GenBank accession number |
|---|---|---|---|---|---|---|---|---|---|---|
| Zt_001 | WAI1822 | Tasmania | Australia | 2014 | Australian National University | ND* | ND | A13 | I14 | MT856842 |
| Zt_002 | WAI2060 | Tasmania | Australia | 2014 | Australian National University | ND | ND | A13 | I14 | MT856842 |
| Zt_003 | WAI1882 | Tasmania | Australia | 2014 | Australian National University | ND | ND | A03 | I02 | MT856832 |
| Zt_004 | WAI1919 | Tasmania | Australia | 2014 | Australian National University | ND | ND | A03 | I02 | MT856832 |
| Zt_005 | WAI1965 | Tasmania | Australia | 2014 | Australian National University | ND | ND | A03 | I02 | MT856832 |
| Zt_006 | WAI1966 | Tasmania | Australia | 2014 | Australian National University | ND | ND | A03 | I02 | MT856832 |
| Zt_007 | WAI1993 | Tasmania | Australia | 2014 | Australian National University | ND | ND | A03 | I02 | MT856832 |
| Zt_008 | WAI2228 | Tasmania | Australia | 2014 | Australian National University | ND | ND | A03 | I02 | MT856832 |
| Zt_009 | WAI2230 | Tasmania | Australia | 2014 | Australian National University | ND | ND | A03 | I02 | MT856832 |
| Zt_010 | WAI2232 | Tasmania | Australia | 2014 | Australian National University | ND | ND | A03 | I02 | MT856832 |
| Zt_011 | WAI1820 | Tasmania | Australia | 2014 | Australian National University | ND | ND | A02 | I02 | MT856831 |
| Zt_012 | WAI1848 | Tasmania | Australia | 2014 | Australian National University | ND | ND | A02 | I02 | MT856831 |
| Zt_013 | WAI1849 | Tasmania | Australia | 2014 | Australian National University | ND | ND | A02 | I02 | MT856831 |
| Zt_014 | WAI1850 | Tasmania | Australia | 2014 | Australian National University | ND | ND | A02 | I02 | MT856831 |
| Zt_015 | WAI1851 | Tasmania | Australia | 2014 | Australian National University | ND | ND | A02 | I02 | MT856831 |
| Zt_016 | WAI1852 | Tasmania | Australia | 2014 | Australian National University | ND | ND | A02 | I02 | MT856831 |
| Zt_017 | WAI1858 | Tasmania | Australia | 2014 | Australian National University | ND | ND | A02 | I02 | MT856831 |
| Zt_018 | WAI1859 | Tasmania | Australia | 2014 | Australian National University | ND | ND | A02 | I02 | MT856831 |
| Zt_019 | WAI1875 | Tasmania | Australia | 2014 | Australian National University | ND | ND | A02 | I02 | MT856831 |
| Zt_020 | WAI1876 | Tasmania | Australia | 2014 | Australian National University | ND | ND | A02 | I02 | MT856831 |
| Zt_021 | WAI1877 | Tasmania | Australia | 2014 | Australian National University | ND | ND | A02 | I02 | MT856831 |
| Zt_022 | WAI1878 | Tasmania | Australia | 2014 | Australian National University | ND | ND | A02 | I02 | MT856831 |
| Zt_023 | WAI1879 | Tasmania | Australia | 2014 | Australian National University | ND | ND | A02 | I02 | MT856831 |
| Zt_024 | WAI1880 | Tasmania | Australia | 2014 | Australian National University | ND | ND | A02 | I02 | MT856831 |
| Zt_025 | WAI1892 | Tasmania | Australia | 2014 | Australian National University | ND | ND | A02 | I02 | MT856831 |
| Zt_026 | WAI1895 | Tasmania | Australia | 2014 | Australian National University | ND | ND | A02 | I02 | MT856831 |
| Zt_027 | WAI1901 | Tasmania | Australia | 2014 | Australian National University | ND | ND | A02 | I02 | MT856831 |
| Zt_028 | WAI1903 | Tasmania | Australia | 2014 | Australian National University | ND | ND | A02 | I02 | MT856831 |
| Zt_029 | WAI1904 | Tasmania | Australia | 2014 | Australian National University | ND | ND | A02 | I02 | MT856831 |
| Zt_030 | WAI1922 | Tasmania | Australia | 2014 | Australian National University | ND | ND | A02 | I02 | MT856831 |
| Zt_031 | WAI1939 | Tasmania | Australia | 2014 | Australian National University | ND | ND | A02 | I02 | MT856831 |
| Zt_032 | WAI1941 | Tasmania | Australia | 2014 | Australian National University | ND | ND | A02 | I02 | MT856831 |
| Zt_033 | WAI1955 | Tasmania | Australia | 2014 | Australian National University | ND | ND | A02 | I02 | MT856831 |
| Zt_034 | WAI1957 | Tasmania | Australia | 2014 | Australian National University | ND | ND | A02 | I02 | MT856831 |
| Zt_035 | WAI1967 | Tasmania | Australia | 2014 | Australian National University | ND | ND | A02 | I02 | MT856831 |
| Zt_036 | WAI1968 | Tasmania | Australia | 2014 | Australian National University | ND | ND | A02 | I02 | MT856831 |
| Zt_037 | WAI1969 | Tasmania | Australia | 2014 | Australian National University | ND | ND | A02 | I02 | MT856831 |
| Zt_038 | WAI1970 | Tasmania | Australia | 2014 | Australian National University | ND | ND | A02 | I02 | MT856831 |
| Zt_039 | WAI1970D | Tasmania | Australia | 2014 | Australian National University | ND | ND | A02 | I02 | MT856831 |
| Zt_040 | WAI1971 | Tasmania | Australia | 2014 | Australian National University | ND | ND | A02 | I02 | MT856831 |
| Zt_041 | WAI1972 | Tasmania | Australia | 2014 | Australian National University | ND | ND | A02 | I02 | MT856831 |
| Zt_042 | WAI1998 | Tasmania | Australia | 2014 | Australian National University | ND | ND | A02 | I02 | MT856831 |
| Zt_043 | WAI2000 | Tasmania | Australia | 2014 | Australian National University | ND | ND | A02 | I02 | MT856831 |
| Zt_044 | WAI2029 | Tasmania | Australia | 2014 | Australian National University | ND | ND | A02 | I02 | MT856831 |
| Zt_045 | WAI2031 | Tasmania | Australia | 2014 | Australian National University | ND | ND | A02 | I02 | MT856831 |
| Zt_046 | WAI2045 | Tasmania | Australia | 2014 | Australian National University | ND | ND | A02 | I02 | MT856831 |
| Zt_047 | WAI2047 | Tasmania | Australia | 2014 | Australian National University | ND | ND | A02 | I02 | MT856831 |
| Zt_048 | WAI2059 | Tasmania | Australia | 2014 | Australian National University | ND | ND | A02 | I02 | MT856831 |
| Zt_049 | WAI2061 | Tasmania | Australia | 2014 | Australian National University | ND | ND | A02 | I02 | MT856831 |
| Zt_050 | WAI2073 | Tasmania | Australia | 2014 | Australian National University | ND | ND | A02 | I02 | MT856831 |
| Zt_051 | WAI2077 | Tasmania | Australia | 2014 | Australian National University | ND | ND | A02 | I02 | MT856831 |
| Zt_052 | WAI2203 | Tasmania | Australia | 2014 | Australian National University | ND | ND | A02 | I02 | MT856831 |
| Zt_053 | WAI2204 | Tasmania | Australia | 2014 | Australian National University | ND | ND | A02 | I02 | MT856831 |
| Zt_054 | WAI2206 | Tasmania | Australia | 2014 | Australian National University | ND | ND | A02 | I02 | MT856831 |
| Zt_055 | WAI2208 | Tasmania | Australia | 2014 | Australian National University | ND | ND | A02 | I02 | MT856831 |
| Zt_056 | WAI2216 | Tasmania | Australia | 2014 | Australian National University | ND | ND | A02 | I02 | MT856831 |
| Zt_057 | WAI2210 | Tasmania | Australia | 2014 | Australian National University | ND | ND | A15 | I02 | MT856844 |
| Zt_058 | WAI2226 | Tasmania | Australia | 2014 | Australian National University | ND | ND | A15 | I02 | MT856844 |
| Zt_059 | CHI 10A | South America | Chile | 2016 | Rothamsted Research | Crac-baer | ND | A06 | I02 | MT856835 |
| Zt_060 | CHI 19B | South America | Chile | 2016 | Rothamsted Research | Crac-baer | ND | A06 | I02 | MT856835 |
| Zt_061 | CHI 2A | South America | Chile | 2016 | Rothamsted Research | Crac-baer | ND | A06 | I02 | MT856835 |
| Zt_062 | CHI 38A | South America | Chile | 2016 | Rothamsted Research | Crac-baer | ND | A06 | I02 | MT856835 |
| Zt_063 | CHI 5A | South America | Chile | 2016 | Rothamsted Research | Crac-baer | ND | A06 | I02 | MT856835 |
| Zt_064 | CHI 7A | South America | Chile | 2016 | Rothamsted Research | Crac-baer | ND | A06 | I02 | MT856835 |
| Zt_065 | CHI 6A | South America | Chile | 2016 | Rothamsted Research | Crac-baer | ND | A03 | I02 | MT856832 |
| Zt_066 | R50 16 | South America | Uruguay | 2016 | Rothamsted Research | Genesis INIA 2375 | 3 (S) | A02 | I02 | MT856831 |
| Zt_067 | R50 27 | South America | Uruguay | 2016 | Rothamsted Research | Genesis INIA 2375 | 3 (S) | A34 | I29 | MT856861 |
| Zt_068 | R50 6 | South America | Uruguay | 2016 | Rothamsted Research | Genesis INIA 2375 | 3 (S) | A02 | I02 | MT856831 |
| Zt_069 | CHI 18B | South America | Chile | 2016 | Rothamsted Research | Crac-baer | ND | A02 | I02 | MT856831 |
| Zt_070 | ER 27A | South America | Argentina | 2016 | Rothamsted Research | DM Fuste | ND | A02 | I02 | MT856831 |
| Zt_071 | ER 30A | South America | Argentina | 2016 | Rothamsted Research | DM Fuste | ND | A02 | I02 | MT856831 |
| Zt_072 | CHI 26A | South America | Chile | 2016 | Rothamsted Research | Crac-baer | ND | A36 | I37 | MT856863 |
| Zt_073 | ER 1A | South America | Argentina | 2016 | Rothamsted Research | DM Fuste | ND | A47 | I03 | MT856873 |
| Zt_074 | R50 12A | South America | Uruguay | 2016 | Rothamsted Research | Genesis INIA 2375 | 3 (S) | A04 | I03 | MT856833 |
| Zt_075 | R50 19 | South America | Uruguay | 2016 | Rothamsted Research | Genesis INIA 2375 | 3 (S) | A04 | I03 | MT856833 |
| Zt_076 | R50 1A | South America | Uruguay | 2016 | Rothamsted Research | Genesis INIA 2375 | 3 (S) | A04 | I03 | MT856833 |
| Zt_077 | R50 20B | South America | Uruguay | 2016 | Rothamsted Research | Genesis INIA 2375 | 3 (S) | A04 | I03 | MT856833 |
| Zt_078 | R50 45A | South America | Uruguay | 2016 | Rothamsted Research | Genesis INIA 2375 | 3 (S) | A04 | I03 | MT856833 |
| Zt_079 | R50 47C | South America | Uruguay | 2016 | Rothamsted Research | Genesis INIA 2375 | 3 (S) | A04 | I03 | MT856833 |
| Zt_080 | R50 8C | South America | Uruguay | 2016 | Rothamsted Research | Genesis INIA 2375 | 3 (S) | A04 | I03 | MT856833 |
| Zt_081 | CHI 36B | South America | Chile | 2016 | Rothamsted Research | Crac-baer | ND | A04 | I03 | MT856833 |

| | | | | | | | | | | |
|---|---|---|---|---|---|---|---|---|---|---|
| Zt_082 | ER 13A | South America | Argentina | 2016 | Rothamsted Research | DM Fuste | ND | A04 | I03 | MT856833 |
| Zt_083 | ER 18A | South America | Argentina | 2016 | Rothamsted Research | DM Fuste | ND | A04 | I03 | MT856833 |
| Zt_084 | ER 56A | South America | Argentina | 2016 | Rothamsted Research | DM Fuste | ND | A04 | I03 | MT856833 |
| Zt_085 | ER 57A | South America | Argentina | 2016 | Rothamsted Research | DM Fuste | ND | A04 | I03 | MT856833 |
| Zt_086 | ER 58A | South America | Argentina | 2016 | Rothamsted Research | DM Fuste | ND | A04 | I03 | MT856833 |
| Zt_087 | ER 61A | South America | Argentina | 2016 | Rothamsted Research | DM Fuste | ND | A04 | I03 | MT856833 |
| Zt_088 | ER 65A | South America | Argentina | 2016 | Rothamsted Research | DM Fuste | ND | A04 | I03 | MT856833 |
| Zt_089 | 11.8 | Mediterranean | Turkey | 2013-2016 | Sirnak University | ND | ND | A24 | I41 | - |
| Zt_090 | 9.2 | Mediterranean | Turkey | 2013-2016 | Sirnak University | ND | ND | A25 | I28 | MT856852 |
| Zt_091 | 3.3 | Mediterranean | Turkey | 2013-2016 | Sirnak University | ND | ND | A27 | I30 | MT856854 |
| Zt_092 | 12.5 | Mediterranean | Turkey | 2013-2016 | Sirnak University | ND | ND | A28 | I34 | MT856855 |
| Zt_093 | 11.5 | Mediterranean | Turkey | 2013-2016 | Sirnak University | ND | ND | A29 | I31 | MT856856 |
| Zt_094 | 3.1 | Mediterranean | Turkey | 2013-2016 | Sirnak University | ND | ND | A14 | I15 | MT856843 |
| Zt_095 | 3.2 | Mediterranean | Turkey | 2013-2016 | Sirnak University | ND | ND | A14 | I15 | MT856843 |
| Zt_096 | 12.1 | Mediterranean | Turkey | 2013-2016 | Sirnak University | ND | ND | A32 | I26 | MT856859 |
| Zt_097 | 10.5 | Mediterranean | Turkey | 2013-2016 | Sirnak University | ND | ND | A17 | I12 | - |
| Zt_098 | 10.6 | Mediterranean | Turkey | 2013-2016 | Sirnak University | ND | ND | A17 | I12 | - |
| Zt_099 | 13.8 | Mediterranean | Turkey | 2013-2016 | Sirnak University | ND | ND | A38 | I23 | MT856865 |
| Zt_100 | 11.4 | Mediterranean | Turkey | 2013-2016 | Sirnak University | ND | ND | A39 | I19 | MT856866 |
| Zt_101 | 12.7 | Mediterranean | Turkey | 2013-2016 | Sirnak University | ND | ND | A40 | I22 | MT856867 |
| Zt_102 | 6.6 | Mediterranean | Turkey | 2013-2016 | Sirnak University | ND | ND | A44 | I25 | MT856871 |
| Zt_103 | 6.3 | Mediterranean | Turkey | 2013-2016 | Sirnak University | ND | ND | A45 | I06 | MT856872 |
| Zt_104 | 6.4 | Mediterranean | Turkey | 2013-2016 | Sirnak University | ND | ND | A46 | I42 | - |
| Zt_105 | 7.1 | Mediterranean | Turkey | 2013-2016 | Sirnak University | ND | ND | A11 | I06 | MT856840 |
| Zt_106 | 10.3 | Mediterranean | Turkey | 2013-2016 | Sirnak University | ND | ND | A11 | I06 | MT856840 |
| Zt_107 | 13.7 | Mediterranean | Turkey | 2013-2016 | Sirnak University | ND | ND | A11 | I06 | MT856840 |
| Zt_108 | 6.1 | Mediterranean | Turkey | 2013-2016 | Sirnak University | ND | ND | A18 | I06 | MT856846 |
| Zt_109 | 7.4 | Mediterranean | Turkey | 2013-2016 | Sirnak University | ND | ND | A18 | I06 | MT856846 |
| Zt_110 | 7.6 | Mediterranean | Turkey | 2013-2016 | Sirnak University | ND | ND | A19 | I16 | MT856847 |
| Zt_111 | 8.7 | Mediterranean | Turkey | 2013-2016 | Sirnak University | ND | ND | A19 | I16 | MT856847 |
| Zt_112 | 3.8 | Mediterranean | Turkey | 2013-2016 | Sirnak University | ND | ND | A04 | I03 | MT856833 |
| Zt_113 | 6.7 | Mediterranean | Turkey | 2013-2016 | Sirnak University | ND | ND | A04 | I03 | MT856833 |
| Zt_114 | 9.4 | Mediterranean | Turkey | 2013-2016 | Sirnak University | ND | ND | A04 | I03 | MT856833 |
| Zt_115 | 9.5 | Mediterranean | Turkey | 2013-2016 | Sirnak University | ND | ND | A04 | I03 | MT856833 |
| Zt_116 | 13.2 | Mediterranean | Turkey | 2013-2016 | Sirnak University | ND | ND | A04 | I03 | MT856833 |
| Zt_117 | 13.4 | Mediterranean | Turkey | 2013-2016 | Sirnak University | ND | ND | A04 | I03 | MT856833 |
| Zt_118 | 3.6 | Mediterranean | Turkey | 2013-2016 | Sirnak University | ND | ND | A20 | I18 | MT856848 |
| Zt_119 | 3.7 | Mediterranean | Turkey | 2013-2016 | Sirnak University | ND | ND | A20 | I18 | MT856848 |
| Zt_120 | 12.2 | Mediterranean | Turkey | 2013-2016 | Sirnak University | ND | ND | A48 | I24 | MT856874 |
| Zt_121 | 6.2 | Mediterranean | Turkey | 2013-2016 | Sirnak University | ND | ND | A05 | I04 | MT856834 |
| Zt_122 | 7.2 | Mediterranean | Turkey | 2013-2016 | Sirnak University | ND | ND | A05 | I04 | MT856834 |
| Zt_123 | 7.3 | Mediterranean | Turkey | 2013-2016 | Sirnak University | ND | ND | A05 | I04 | MT856834 |
| Zt_124 | 8.1 | Mediterranean | Turkey | 2013-2016 | Sirnak University | ND | ND | A05 | I04 | MT856834 |
| Zt_125 | 8.4 | Mediterranean | Turkey | 2013-2016 | Sirnak University | ND | ND | A05 | I04 | MT856834 |
| Zt_126 | 8.6 | Mediterranean | Turkey | 2013-2016 | Sirnak University | ND | ND | A05 | I04 | MT856834 |
| Zt_127 | 8.8 | Mediterranean | Turkey | 2013-2016 | Sirnak University | ND | ND | A05 | I04 | MT856834 |
| Zt_128 | 12.4 | Mediterranean | Turkey | 2013-2016 | Sirnak University | ND | ND | A05 | I04 | MT856834 |
| Zt_129 | 8.5 | Mediterranean | Turkey | 2013-2016 | Sirnak University | ND | ND | A50 | I32 | MT856876 |
| Zt_130 | 10.4 | Mediterranean | Turkey | 2013-2016 | Sirnak University | ND | ND | A51 | I43 | MT856877 |
| Zt_131 | 7.5 | Mediterranean | Turkey | 2013-2016 | Sirnak University | ND | ND | A09 | I08 | MT856838 |
| Zt_132 | 7.8 | Mediterranean | Turkey | 2013-2016 | Sirnak University | ND | ND | A09 | I08 | MT856838 |
| Zt_133 | 9.3 | Mediterranean | Turkey | 2013-2016 | Sirnak University | ND | ND | A09 | I08 | MT856838 |
| Zt_134 | 10.1 | Mediterranean | Turkey | 2013-2016 | Sirnak University | ND | ND | A09 | I08 | MT856838 |
| Zt_135 | 6.5 | Mediterranean | Turkey | 2013-2016 | Sirnak University | ND | ND | A37 | I40 | MT856864 |
| Zt_136 | 6.8 | Mediterranean | Turkey | 2013-2016 | Sirnak University | ND | ND | A10 | I09 | MT856839 |
| Zt_137 | 9.1 | Mediterranean | Turkey | 2013-2016 | Sirnak University | ND | ND | A10 | I09 | MT856839 |
| Zt_138 | 10.2 | Mediterranean | Turkey | 2013-2016 | Sirnak University | ND | ND | A10 | I09 | MT856839 |
| Zt_139 | 3.4 | Mediterranean | Turkey | 2013-2016 | Sirnak University | ND | ND | A10 | I09 | MT856839 |
| Zt_140 | 7.7 | Mediterranean | Turkey | 2013-2016 | Sirnak University | ND | ND | A12 | I11 | MT856841 |
| Zt_141 | 8.3 | Mediterranean | Turkey | 2013-2016 | Sirnak University | ND | ND | A12 | I11 | MT856841 |
| Zt_142 | 9.6 | Mediterranean | Turkey | 2013-2016 | Sirnak University | ND | ND | A12 | I11 | MT856841 |
| Zt_143 | 8.2 | Mediterranean | Turkey | 2013-2016 | Sirnak University | ND | ND | A22 | I10 | MT856850 |
| Zt_144 | 9.8 | Mediterranean | Turkey | 2013-2016 | Sirnak University | ND | ND | A22 | I10 | MT856850 |
| Zt_145 | 9.7 | Mediterranean | Turkey | 2013-2016 | Sirnak University | ND | ND | A23 | I10 | MT856851 |
| Zt_146 | Ore 52 | Oregon | USA | 2016 | Rothamsted Research | Kaseberg | 7 (S) | A31 | I39 | MT856858 |
| Zt_147 | Ore 1 | Oregon | USA | 2016 | Rothamsted Research | Kaseberg | 7 (S) | A02 | I02 | MT856831 |
| Zt_148 | Ore 10 | Oregon | USA | 2016 | Rothamsted Research | Kaseberg | 7 (S) | A02 | I02 | MT856831 |
| Zt_149 | Ore 12 | Oregon | USA | 2016 | Rothamsted Research | Kaseberg | 7 (S) | A02 | I02 | MT856831 |
| Zt_150 | Ore 13 | Oregon | USA | 2016 | Rothamsted Research | Kaseberg | 7 (S) | A02 | I02 | MT856831 |
| Zt_151 | Ore 14 | Oregon | USA | 2016 | Rothamsted Research | Kaseberg | 7 (S) | A02 | I02 | MT856831 |
| Zt_152 | Ore 16 | Oregon | USA | 2016 | Rothamsted Research | Kaseberg | 7 (S) | A02 | I02 | MT856831 |
| Zt_153 | Ore 17 | Oregon | USA | 2016 | Rothamsted Research | Kaseberg | 7 (S) | A02 | I02 | MT856831 |
| Zt_154 | Ore 19 | Oregon | USA | 2016 | Rothamsted Research | Kaseberg | 7 (S) | A02 | I02 | MT856831 |
| Zt_155 | Ore 2 | Oregon | USA | 2016 | Rothamsted Research | Kaseberg | 7 (S) | A02 | I02 | MT856831 |
| Zt_156 | Ore 20 | Oregon | USA | 2016 | Rothamsted Research | Kaseberg | 7 (S) | A02 | I02 | MT856831 |
| Zt_157 | Ore 21 | Oregon | USA | 2016 | Rothamsted Research | Kaseberg | 7 (S) | A02 | I02 | MT856831 |
| Zt_158 | Ore 22 | Oregon | USA | 2016 | Rothamsted Research | Kaseberg | 7 (S) | A02 | I02 | MT856831 |
| Zt_159 | Ore 24 | Oregon | USA | 2016 | Rothamsted Research | Kaseberg | 7 (S) | A02 | I02 | MT856831 |
| Zt_160 | Ore 25 | Oregon | USA | 2016 | Rothamsted Research | Kaseberg | 7 (S) | A02 | I02 | MT856831 |
| Zt_161 | Ore 26 | Oregon | USA | 2016 | Rothamsted Research | Kaseberg | 7 (S) | A02 | I02 | MT856831 |
| Zt_162 | Ore 27 | Oregon | USA | 2016 | Rothamsted Research | Kaseberg | 7 (S) | A02 | I02 | MT856831 |
| Zt_163 | Ore 3 | Oregon | USA | 2016 | Rothamsted Research | Kaseberg | 7 (S) | A02 | I02 | MT856831 |
| Zt_164 | Ore 30 | Oregon | USA | 2016 | Rothamsted Research | Kaseberg | 7 (S) | A02 | I02 | MT856831 |
| Zt_165 | Ore 31 | Oregon | USA | 2016 | Rothamsted Research | Kaseberg | 7 (S) | A02 | I02 | MT856831 |
| Zt_166 | Ore 33 | Oregon | USA | 2016 | Rothamsted Research | Kaseberg | 7 (S) | A02 | I02 | MT856831 |
| Zt_167 | Ore 34 | Oregon | USA | 2016 | Rothamsted Research | Kaseberg | 7 (S) | A02 | I02 | MT856831 |
| Zt_168 | Ore 36 | Oregon | USA | 2016 | Rothamsted Research | Kaseberg | 7 (S) | A02 | I02 | MT856831 |

| | | | | | | | | | | |
|---|---|---|---|---|---|---|---|---|---|---|
| Zt_169 | Ore 38 | Oregon | USA | 2016 | Rothamsted Research | Kaseberg | 7 (S) | A02 | I02 | MT856831 |
| Zt_170 | Ore 39 | Oregon | USA | 2016 | Rothamsted Research | Kaseberg | 7 (S) | A02 | I02 | MT856831 |
| Zt_171 | Ore 4 | Oregon | USA | 2016 | Rothamsted Research | Kaseberg | 7 (S) | A02 | I02 | MT856831 |
| Zt_172 | Ore 41 | Oregon | USA | 2016 | Rothamsted Research | Kaseberg | 7 (S) | A02 | I02 | MT856831 |
| Zt_173 | Ore 42 | Oregon | USA | 2016 | Rothamsted Research | Kaseberg | 7 (S) | A02 | I02 | MT856831 |
| Zt_174 | Ore 43 | Oregon | USA | 2016 | Rothamsted Research | Kaseberg | 7 (S) | A02 | I02 | MT856831 |
| Zt_175 | Ore 45 | Oregon | USA | 2016 | Rothamsted Research | Kaseberg | 7 (S) | A02 | I02 | MT856831 |
| Zt_176 | Ore 46 | Oregon | USA | 2016 | Rothamsted Research | Kaseberg | 7 (S) | A02 | I02 | MT856831 |
| Zt_177 | Ore 48 | Oregon | USA | 2016 | Rothamsted Research | Kaseberg | 7 (S) | A02 | I02 | MT856831 |
| Zt_178 | Ore 49 | Oregon | USA | 2016 | Rothamsted Research | Kaseberg | 7 (S) | A02 | I02 | MT856831 |
| Zt_179 | Ore 5 | Oregon | USA | 2016 | Rothamsted Research | Kaseberg | 7 (S) | A02 | I02 | MT856831 |
| Zt_180 | Ore 50 | Oregon | USA | 2016 | Rothamsted Research | Kaseberg | 7 (S) | A02 | I02 | MT856831 |
| Zt_181 | Ore 51 | Oregon | USA | 2016 | Rothamsted Research | Kaseberg | 7 (S) | A02 | I02 | MT856831 |
| Zt_182 | Ore 53 | Oregon | USA | 2016 | Rothamsted Research | Kaseberg | 7 (S) | A02 | I02 | MT856831 |
| Zt_183 | Ore 54 | Oregon | USA | 2016 | Rothamsted Research | Kaseberg | 7 (S) | A02 | I02 | MT856831 |
| Zt_184 | Ore 55 | Oregon | USA | 2016 | Rothamsted Research | Kaseberg | 7 (S) | A02 | I02 | MT856831 |
| Zt_185 | Ore 56 | Oregon | USA | 2016 | Rothamsted Research | Kaseberg | 7 (S) | A02 | I02 | MT856831 |
| Zt_186 | Ore 57 | Oregon | USA | 2016 | Rothamsted Research | Kaseberg | 7 (S) | A02 | I02 | MT856831 |
| Zt_187 | Ore 6 | Oregon | USA | 2016 | Rothamsted Research | Kaseberg | 7 (S) | A02 | I02 | MT856831 |
| Zt_188 | Ore 8 | Oregon | USA | 2016 | Rothamsted Research | Kaseberg | 7 (S) | A02 | I02 | MT856831 |
| Zt_189 | Ore 9 | Oregon | USA | 2016 | Rothamsted Research | Kaseberg | 7 (S) | A02 | I02 | MT856831 |
| Zt_190 | Ore 18 | Oregon | USA | 2016 | Rothamsted Research | Kaseberg | 7 (S) | A08 | I07 | MT856837 |
| Zt_191 | Ore 32 | Oregon | USA | 2016 | Rothamsted Research | Kaseberg | 7 (S) | A08 | I07 | MT856837 |
| Zt_192 | Ore 37 | Oregon | USA | 2016 | Rothamsted Research | Kaseberg | 7 (S) | A08 | I07 | MT856837 |
| Zt_193 | Ore 44 | Oregon | USA | 2016 | Rothamsted Research | Kaseberg | 7 (S) | A08 | I07 | MT856837 |
| Zt_194 | Ore 47 | Oregon | USA | 2016 | Rothamsted Research | Kaseberg | 7 (S) | A08 | I07 | MT856837 |
| Zt_195 | Ore 35 | Oregon | USA | 2016 | Rothamsted Research | Kaseberg | 7 (S) | A42 | I20 | MT856869 |
| Zt_196 | Ore 59 | Oregon | USA | 2016 | Rothamsted Research | Kaseberg | 7 (S) | A43 | I38 | MT856870 |
| Zt_197 | Ore 40 | Oregon | USA | 2016 | Rothamsted Research | Kaseberg | 7 (S) | A49 | I35 | MT856875 |
| Zt_198 | 7 | Western Europe | Scotland | 2015 | Rothamsted Research | Consort | 7 (S) | A26 | I27 | MT856853 |
| Zt_199 | NT-S219-101.10 | Western Europe | England | 2015 | Rothamsted Research | KWS Cashel | 3 (S) | A30 | I36 | MT856857 |
| Zt_200 | A2-A2016.30 | Western Europe | England | 2016 | Rothamsted Research | Cougar | 1 (R) | A33 | I02 | MT856860 |
| Zt_201 | BASF.10 | Western Europe | England | 2015 | Rothamsted Research | Cougar | 1 (R) | A03 | I02 | MT856832 |
| Zt_202 | BASF.5 | Western Europe | England | 2015 | Rothamsted Research | Cougar | 1 (R) | A03 | I02 | MT856832 |
| Zt_203 | CRD2015.10 | Western Europe | England | 2015 | Rothamsted Research | Cougar | 1 (R) | A03 | I02 | MT856832 |
| Zt_204 | 11 | Western Europe | England | 2015 | Rothamsted Research | KWS Santiago | 1 (R) | A03 | I02 | MT856832 |
| Zt_205 | 1 | Western Europe | England | 2015 | Rothamsted Research | Consort | 7 (S) | A03 | I02 | MT856832 |
| Zt_206 | CRD.GB23 | Western Europe | England | 2015 | Rothamsted Research | Gallant | 7 (S) | A03 | I02 | MT856832 |
| Zt_207 | CRD2015.G15 | Western Europe | England | 2015 | Rothamsted Research | Gallant | 7 (S) | A03 | I02 | MT856832 |
| Zt_208 | FT1.6 | Western Europe | England | 2015 | Rothamsted Research | Gallant | 7 (S) | A03 | I02 | MT856832 |
| Zt_209 | NTS311 321.2 | Western Europe | England | 2015 | Rothamsted Research | KWS Cashel | 3 (S) | A03 | I02 | MT856832 |
| Zt_210 | NT-S31-239.5 | Western Europe | England | 2015 | Rothamsted Research | KWS Cashel | 3 (S) | A03 | I02 | MT856832 |
| Zt_211 | NT-S319-121.1 | Western Europe | England | 2015 | Rothamsted Research | KWS Cashel | 3 (S) | A03 | I02 | MT856832 |
| Zt_212 | Syn2.6 | Western Europe | France | 2015 | Rothamsted Research | Cellule | ND | A03 | I02 | MT856832 |
| Zt_213 | 15 | Western Europe | Germany | 2015 | Rothamsted Research | Tobac | ND | A03 | I02 | MT856832 |
| Zt_214 | UPL1.6 | Western Europe | Germany | 2015 | Rothamsted Research | Tobac | ND | A03 | I02 | MT856832 |
| Zt_215 | FT1.8 | Western Europe | England | 2015 | Rothamsted Research | Zulu | 1 (R) | A03 | I02 | MT856832 |
| Zt_216 | 020A | Western Europe | England | 2016 | Rothamsted Research | Cougar | 1 (R) | A03 | I02 | MT856832 |
| Zt_217 | M3 2016.27 | Western Europe | England | 2016 | Rothamsted Research | Cordiale | 7 (S) | A03 | I02 | MT856832 |
| Zt_218 | A2-A2016.6 | Western Europe | England | 2016 | Rothamsted Research | Consort | 7 (S) | A03 | I02 | MT856832 |
| Zt_219 | NT1 2016.19 | Western Europe | England | 2016 | Rothamsted Research | Dickens | 1 (R) | A03 | I02 | MT856832 |
| Zt_220 | V7 2016.1 | Western Europe | England | 2016 | Rothamsted Research | Dickens | 1 (R) | A03 | I02 | MT856832 |
| Zt_221 | T1-A 2017.12 | Western Europe | Ireland | 2017 | Rothamsted Research | KWS Lumos | 7 (S) | A03 | I02 | MT856832 |
| Zt_222 | T1-A 2017.39 | Western Europe | Ireland | 2017 | Rothamsted Research | KWS Lumos | 7 (S) | A03 | I02 | MT856832 |
| Zt_223 | S1-B.27 | Western Europe | Scotland | 2017 | Rothamsted Research | ND | ND | A03 | I02 | MT856832 |
| Zt_224 | S1-B.28 | Western Europe | Scotland | 2017 | Rothamsted Research | ND | ND | A03 | I02 | MT856832 |
| Zt_225 | S1-B.31 | Western Europe | Scotland | 2017 | Rothamsted Research | ND | ND | A03 | I02 | MT856832 |
| Zt_226 | 19 | Western Europe | England | 2015 | Rothamsted Research | Cougar | 1 (R) | A02 | I02 | MT856831 |
| Zt_227 | 20 | Western Europe | England | 2015 | Rothamsted Research | Cougar | 1 (R) | A02 | I02 | MT856831 |
| Zt_228 | CRD2015.17 | Western Europe | England | 2015 | Rothamsted Research | Cougar | 1 (R) | A02 | I02 | MT856831 |
| Zt_229 | CRD2015.18 | Western Europe | England | 2015 | Rothamsted Research | Cougar | 1 (R) | A02 | I02 | MT856831 |
| Zt_230 | CRD2015.19 | Western Europe | England | 2015 | Rothamsted Research | Cougar | 1 (R) | A02 | I02 | MT856831 |
| Zt_231 | CRD2015.21 | Western Europe | England | 2015 | Rothamsted Research | Cougar | 1 (R) | A02 | I02 | MT856831 |
| Zt_232 | CRD2015.34 | Western Europe | England | 2015 | Rothamsted Research | Cougar | 1 (R) | A02 | I02 | MT856831 |
| Zt_233 | CRD2015.4 | Western Europe | England | 2015 | Rothamsted Research | Cougar | 1 (R) | A02 | I02 | MT856831 |
| Zt_234 | GS26.41 | Western Europe | England | 2015 | Rothamsted Research | Cougar | 1 (R) | A02 | I02 | MT856831 |
| Zt_235 | GS26.42 | Western Europe | England | 2015 | Rothamsted Research | Cougar | 1 (R) | A02 | I02 | MT856831 |
| Zt_236 | 17 | Western Europe | France | 2015 | Rothamsted Research | Trapez | 1 (R) | A02 | I02 | MT856831 |
| Zt_237 | 10 | Western Europe | England | 2015 | Rothamsted Research | Crusoe | 7 (S) | A02 | I02 | MT856831 |
| Zt_238 | 12 | Western Europe | England | 2015 | Rothamsted Research | JB Diego | 7 (S) | A02 | I02 | MT856831 |
| Zt_239 | 8 | Western Europe | Scotland | 2015 | Rothamsted Research | Consort | 7 (S) | A02 | I02 | MT856831 |
| Zt_240 | 9 | Western Europe | Scotland | 2015 | Rothamsted Research | Consort | 7 (S) | A02 | I02 | MT856831 |
| Zt_241 | R15-46 | Western Europe | England | 2015 | Rothamsted Research | Dickens | 1 (R) | A02 | I02 | MT856831 |
| Zt_242 | 18 | Western Europe | England | 2015 | Rothamsted Research | Gallant | 7 (S) | A02 | I02 | MT856831 |
| Zt_243 | CRD.GB35 | Western Europe | England | 2015 | Rothamsted Research | Gallant | 7 (S) | A02 | I02 | MT856831 |
| Zt_244 | CRD2015.G3 | Western Europe | England | 2015 | Rothamsted Research | Gallant | 7 (S) | A02 | I02 | MT856831 |
| Zt_245 | FT1.17 | Western Europe | England | 2015 | Rothamsted Research | Gallant | 7 (S) | A02 | I02 | MT856831 |
| Zt_246 | 2 | Western Europe | England | 2015 | Rothamsted Research | KWS Cashel | 3 (S) | A02 | I02 | MT856831 |
| Zt_247 | 3 | Western Europe | England | 2015 | Rothamsted Research | KWS Cashel | 3 (S) | A02 | I02 | MT856831 |
| Zt_248 | 5 | Western Europe | England | 2015 | Rothamsted Research | KWS Cashel | 3 (S) | A02 | I02 | MT856831 |
| Zt_249 | 6 | Western Europe | England | 2015 | Rothamsted Research | KWS Cashel | 3 (S) | A35 | I21 | MT856862 |
| Zt_250 | NT15-119.11 | Western Europe | England | 2015 | Rothamsted Research | KWS Cashel | 3 (S) | A02 | I02 | MT856831 |
| Zt_251 | NT15-239.7 | Western Europe | England | 2015 | Rothamsted Research | KWS Cashel | 3 (S) | A02 | I02 | MT856831 |
| Zt_252 | NT15-239.8 | Western Europe | England | 2015 | Rothamsted Research | KWS Cashel | 3 (S) | A02 | I02 | MT856831 |
| Zt_253 | NT-S219-104.3 | Western Europe | England | 2015 | Rothamsted Research | KWS Cashel | 3 (S) | A02 | I02 | MT856831 |
| Zt_254 | NT-S219-211.8 | Western Europe | England | 2015 | Rothamsted Research | KWS Cashel | 3 (S) | A02 | I02 | MT856831 |
| Zt_255 | NT-S311-118.9 | Western Europe | England | 2015 | Rothamsted Research | KWS Cashel | 3 (S) | A02 | I02 | MT856831 |

| | | | | | | | | | | |
|---|---|---|---|---|---|---|---|---|---|---|
| Zt_256 | NT-S319-121.3 | Western Europe | England | 2015 | Rothamsted Research | KWS Cashel | 3 (S) | A02 | I02 | MT856831 |
| Zt_257 | NT-S319-121.4 | Western Europe | England | 2015 | Rothamsted Research | KWS Cashel | 3 (S) | A02 | I02 | MT856831 |
| Zt_258 | NT-S319-317.2 | Western Europe | England | 2015 | Rothamsted Research | KWS Cashel | 3 (S) | A02 | I02 | MT856831 |
| Zt_259 | NT-S319-317.4 | Western Europe | England | 2015 | Rothamsted Research | KWS Cashel | 3 (S) | A02 | I02 | MT856831 |
| Zt_260 | NT-S319-317.5 | Western Europe | England | 2015 | Rothamsted Research | KWS Cashel | 3 (S) | A02 | I02 | MT856831 |
| Zt_261 | Syn2.10 | Western Europe | France | 2015 | Rothamsted Research | Cellule | ND | A02 | I02 | MT856831 |
| Zt_262 | Syn2.11 | Western Europe | France | 2015 | Rothamsted Research | Cellule | ND | A02 | I02 | MT856831 |
| Zt_263 | Syn2.3 | Western Europe | France | 2015 | Rothamsted Research | Cellule | ND | A02 | I02 | MT856831 |
| Zt_264 | Syn2.4 | Western Europe | France | 2015 | Rothamsted Research | Cellule | ND | A02 | I02 | MT856831 |
| Zt_265 | Syn2.5 | Western Europe | France | 2015 | Rothamsted Research | Cellule | ND | A02 | I02 | MT856831 |
| Zt_266 | Syn2.7 | Western Europe | France | 2015 | Rothamsted Research | Cellule | ND | A02 | I02 | MT856831 |
| Zt_267 | 14 | Western Europe | Germany | 2015 | Rothamsted Research | JB Asano | ND | A02 | I02 | MT856831 |
| Zt_268 | UPL1.1 | Western Europe | Germany | 2015 | Rothamsted Research | Tobac | ND | A02 | I02 | MT856831 |
| Zt_269 | UPL1.11 | Western Europe | Germany | 2015 | Rothamsted Research | Tobac | ND | A02 | I02 | MT856831 |
| Zt_270 | UPL1.2 | Western Europe | Germany | 2015 | Rothamsted Research | Tobac | ND | A02 | I02 | MT856831 |
| Zt_271 | UPL1.3 | Western Europe | Germany | 2015 | Rothamsted Research | Tobac | ND | A02 | I02 | MT856831 |
| Zt_272 | UPL1.4 | Western Europe | Germany | 2015 | Rothamsted Research | Tobac | ND | A02 | I02 | MT856831 |
| Zt_273 | UPL1.5V2-R | Western Europe | Germany | 2015 | Rothamsted Research | Tobac | ND | A02 | I02 | MT856831 |
| Zt_274 | UPL1.7 | Western Europe | Germany | 2015 | Rothamsted Research | Tobac | ND | A02 | I02 | MT856831 |
| Zt_275 | UPL1.9 | Western Europe | Germany | 2015 | Rothamsted Research | Tobac | ND | A02 | I02 | MT856831 |
| Zt_276 | 014C | Western Europe | England | 2016 | Rothamsted Research | Cougar | 1 (R) | A02 | I02 | MT856831 |
| Zt_277 | 062A | Western Europe | England | 2016 | Rothamsted Research | Cougar | 1 (R) | A02 | I02 | MT856831 |
| Zt_278 | A2-A2016.47 | Western Europe | England | 2016 | Rothamsted Research | Cougar | 1 (R) | A02 | I02 | MT856831 |
| Zt_279 | A3-A.2 | Western Europe | England | 2016 | Rothamsted Research | Cougar | 1 (R) | A02 | I02 | MT856831 |
| Zt_280 | A3-A.36 | Western Europe | England | 2016 | Rothamsted Research | Cougar | 1 (R) | A02 | I02 | MT856831 |
| Zt_281 | A3-A.37 | Western Europe | England | 2016 | Rothamsted Research | Cougar | 1 (R) | A02 | I02 | MT856831 |
| Zt_282 | V1.12 | Western Europe | England | 2016 | Rothamsted Research | Evolution | 1 (R) | A02 | I02 | MT856831 |
| Zt_283 | V1.16 | Western Europe | England | 2016 | Rothamsted Research | Evolution | 1 (R) | A02 | I02 | MT856831 |
| Zt_284 | V1.31 | Western Europe | England | 2016 | Rothamsted Research | Evolution | 1 (R) | A02 | I02 | MT856831 |
| Zt_285 | V1.45 | Western Europe | England | 2016 | Rothamsted Research | Evolution | 1 (R) | A02 | I02 | MT856831 |
| Zt_286 | V4 2016.31 | Western Europe | England | 2016 | Rothamsted Research | Evolution | 1 (R) | A02 | I02 | MT856831 |
| Zt_287 | V4.35 | Western Europe | England | 2016 | Rothamsted Research | Evolution | 1 (R) | A02 | I02 | MT856831 |
| Zt_288 | 009A | Western Europe | England | 2016 | Rothamsted Research | KWS Santiago | 1 (R) | A02 | I02 | MT856831 |
| Zt_289 | M384 | Western Europe | England | 2016 | Rothamsted Research | KWS Santiago | 1 (R) | A02 | I02 | MT856831 |
| Zt_290 | M389 | Western Europe | England | 2016 | Rothamsted Research | KWS Santiago | 1 (R) | A02 | I02 | MT856831 |
| Zt_291 | M4 2016.42 | Western Europe | England | 2016 | Rothamsted Research | Alchemy | 7 (S) | A02 | I02 | MT856831 |
| Zt_292 | M3 2016.23 | Western Europe | England | 2016 | Rothamsted Research | Cordiale | 7 (S) | A02 | I02 | MT856831 |
| Zt_293 | M3 2016.42 | Western Europe | England | 2016 | Rothamsted Research | Cordiale | 7 (S) | A02 | I02 | MT856831 |
| Zt_294 | M3.1 | Western Europe | England | 2016 | Rothamsted Research | Cordiale | 7 (S) | A02 | I02 | MT856831 |
| Zt_295 | M3.11 | Western Europe | England | 2016 | Rothamsted Research | Cordiale | 7 (S) | A02 | I02 | MT856831 |
| Zt_296 | M3.12 | Western Europe | England | 2016 | Rothamsted Research | Cordiale | 7 (S) | A02 | I02 | MT856831 |
| Zt_297 | M3.13 | Western Europe | England | 2016 | Rothamsted Research | Cordiale | 7 (S) | A02 | I02 | MT856831 |
| Zt_298 | M3.2 | Western Europe | England | 2016 | Rothamsted Research | Cordiale | 7 (S) | A02 | I02 | MT856831 |
| Zt_299 | M3.23 | Western Europe | England | 2016 | Rothamsted Research | Cordiale | 7 (S) | A02 | I02 | MT856831 |
| Zt_300 | M3.25 | Western Europe | England | 2016 | Rothamsted Research | Cordiale | 7 (S) | A02 | I02 | MT856831 |
| Zt_301 | M9 2016.26 | Western Europe | England | 2016 | Rothamsted Research | Cordiale | 7 (S) | A02 | I02 | MT856831 |
| Zt_302 | M9 2016.4 | Western Europe | England | 2016 | Rothamsted Research | Cordiale | 7 (S) | A02 | I02 | MT856831 |
| Zt_303 | V4 2016.21 | Western Europe | England | 2016 | Rothamsted Research | JB Diego | 7 (S) | A02 | I02 | MT856831 |
| Zt_304 | R16.1 | Western Europe | England | 2016 | Rothamsted Research | Reflection | 7 (S) | A02 | I02 | MT856831 |
| Zt_305 | R16.18 | Western Europe | England | 2016 | Rothamsted Research | Reflection | 7 (S) | A02 | I02 | MT856831 |
| Zt_306 | R16.19 | Western Europe | England | 2016 | Rothamsted Research | Reflection | 7 (S) | A02 | I02 | MT856831 |
| Zt_307 | R16.23 | Western Europe | England | 2016 | Rothamsted Research | Reflection | 7 (S) | A02 | I02 | MT856831 |
| Zt_308 | R16.37 | Western Europe | England | 2016 | Rothamsted Research | Reflection | 7 (S) | A02 | I02 | MT856831 |
| Zt_309 | A2-B.11 | Western Europe | England | 2016 | Rothamsted Research | Consort | 7 (S) | A02 | I02 | MT856831 |
| Zt_310 | A2-B.21 | Western Europe | England | 2016 | Rothamsted Research | Consort | 7 (S) | A02 | I02 | MT856831 |
| Zt_311 | A3-A.3 | Western Europe | England | 2016 | Rothamsted Research | Consort | 7 (S) | A02 | I02 | MT856831 |
| Zt_312 | A3-B.16 | Western Europe | England | 2016 | Rothamsted Research | Consort | 7 (S) | A02 | I02 | MT856831 |
| Zt_313 | A3-B.3 | Western Europe | England | 2016 | Rothamsted Research | Consort | 7 (S) | A02 | I02 | MT856831 |
| Zt_314 | A3-C.11 | Western Europe | England | 2016 | Rothamsted Research | Consort | 7 (S) | A02 | I02 | MT856831 |
| Zt_315 | A3-C.12 | Western Europe | England | 2016 | Rothamsted Research | Consort | 7 (S) | A02 | I02 | MT856831 |
| Zt_316 | A3-C.17 | Western Europe | England | 2016 | Rothamsted Research | Consort | 7 (S) | A02 | I02 | MT856831 |
| Zt_317 | A3-C.18 | Western Europe | England | 2016 | Rothamsted Research | Consort | 7 (S) | A02 | I02 | MT856831 |
| Zt_318 | S2-C.13 | Western Europe | Scotland | 2016 | Rothamsted Research | Consort | 7 (S) | A02 | I02 | MT856831 |
| Zt_319 | S3-A.3 | Western Europe | Scotland | 2016 | Rothamsted Research | Consort | 7 (S) | A02 | I02 | MT856831 |
| Zt_320 | S3-A.9 | Western Europe | Scotland | 2016 | Rothamsted Research | Consort | 7 (S) | A02 | I02 | MT856831 |
| Zt_321 | M31 10 | Western Europe | England | 2016 | Rothamsted Research | Dickens | 1 (R) | A02 | I02 | MT856831 |
| Zt_322 | M31 13 | Western Europe | England | 2016 | Rothamsted Research | Dickens | 1 (R) | A02 | I02 | MT856831 |
| Zt_323 | M31 16 | Western Europe | England | 2016 | Rothamsted Research | Dickens | 1 (R) | A02 | I02 | MT856831 |
| Zt_324 | M38 19 | Western Europe | England | 2016 | Rothamsted Research | Dickens | 1 (R) | A02 | I02 | MT856831 |
| Zt_325 | M38 27 | Western Europe | England | 2016 | Rothamsted Research | Dickens | 1 (R) | A02 | I02 | MT856831 |
| Zt_326 | NT1 2016.1 | Western Europe | England | 2016 | Rothamsted Research | Dickens | 1 (R) | A02 | I02 | MT856831 |
| Zt_327 | NT1 2016.21 | Western Europe | England | 2016 | Rothamsted Research | Dickens | 1 (R) | A02 | I02 | MT856831 |
| Zt_328 | NT1 2016.9 | Western Europe | England | 2016 | Rothamsted Research | Dickens | 1 (R) | A02 | I02 | MT856831 |
| Zt_329 | NT2-B.1 | Western Europe | England | 2016 | Rothamsted Research | Dickens | 1 (R) | A02 | I02 | MT856831 |
| Zt_330 | NT2-B.10 | Western Europe | England | 2016 | Rothamsted Research | Dickens | 1 (R) | A02 | I02 | MT856831 |
| Zt_331 | NT2-C.14 | Western Europe | England | 2016 | Rothamsted Research | Dickens | 1 (R) | A02 | I02 | MT856831 |
| Zt_332 | NT3-B.1 | Western Europe | England | 2016 | Rothamsted Research | Dickens | 1 (R) | A02 | I02 | MT856831 |
| Zt_333 | NT3-C.13 | Western Europe | England | 2016 | Rothamsted Research | Dickens | 1 (R) | A02 | I02 | MT856831 |
| Zt_334 | NT3-C.9 | Western Europe | England | 2016 | Rothamsted Research | Dickens | 1 (R) | A02 | I02 | MT856831 |
| Zt_335 | NT4-A.1 | Western Europe | England | 2016 | Rothamsted Research | Dickens | 1 (R) | A02 | I02 | MT856831 |
| Zt_336 | NT4-A.10 | Western Europe | England | 2016 | Rothamsted Research | Dickens | 1 (R) | A02 | I02 | MT856831 |
| Zt_337 | NT4-A.2 | Western Europe | England | 2016 | Rothamsted Research | Dickens | 1 (R) | A02 | I02 | MT856831 |
| Zt_338 | NT4-B.9 | Western Europe | England | 2016 | Rothamsted Research | Dickens | 1 (R) | A02 | I02 | MT856831 |
| Zt_339 | NT4-C.11 | Western Europe | England | 2016 | Rothamsted Research | Dickens | 1 (R) | A02 | I02 | MT856831 |
| Zt_340 | 069C | Western Europe | England | 2016 | Rothamsted Research | KWS Cashel | 3 (S) | A02 | I02 | MT856831 |
| Zt_341 | 001B | Western Europe | England | 2016 | Rothamsted Research | Marston | 1 (S) | A02 | I02 | MT856831 |
| Zt_342 | V5 2016.37 | Western Europe | England | 2016 | Rothamsted Research | Zulu | 1 (R) | A02 | I02 | MT856831 |

| | | | | | | | | | | |
|---|---|---|---|---|---|---|---|---|---|---|
| Zt_343 | M3 2016.10 | Western Europe | ND | 2016 | Rothamsted Research | ND | ND | A02 | I02 | MT856831 |
| Zt_344 | M3 2016.35 | Western Europe | ND | 2016 | Rothamsted Research | ND | ND | A02 | I02 | MT856831 |
| Zt_345 | RR17.1 | Western Europe | England | 2017 | Rothamsted Research | KWS Siskin | 1 (R) | A02 | I02 | MT856831 |
| Zt_346 | RR17.10 | Western Europe | England | 2017 | Rothamsted Research | KWS Siskin | 1 (R) | A02 | I02 | MT856831 |
| Zt_347 | RR17.11 | Western Europe | England | 2017 | Rothamsted Research | KWS Siskin | 1 (R) | A02 | I02 | MT856831 |
| Zt_348 | RR17.12 | Western Europe | England | 2017 | Rothamsted Research | KWS Siskin | 1 (R) | A02 | I02 | MT856831 |
| Zt_349 | RR17.13 | Western Europe | England | 2017 | Rothamsted Research | KWS Siskin | 1 (R) | A02 | I02 | MT856831 |
| Zt_350 | RR17.14 | Western Europe | England | 2017 | Rothamsted Research | KWS Siskin | 1 (R) | A02 | I02 | MT856831 |
| Zt_351 | RR17.15 | Western Europe | England | 2017 | Rothamsted Research | KWS Siskin | 1 (R) | A02 | I02 | MT856831 |
| Zt_352 | RR17.3 | Western Europe | England | 2017 | Rothamsted Research | KWS Siskin | 1 (R) | A02 | I02 | MT856831 |
| Zt_353 | RR17.4 | Western Europe | England | 2017 | Rothamsted Research | KWS Siskin | 1 (R) | A02 | I02 | MT856831 |
| Zt_354 | RR17.5 | Western Europe | England | 2017 | Rothamsted Research | KWS Siskin | 1 (R) | A02 | I02 | MT856831 |
| Zt_355 | RR17.6 | Western Europe | England | 2017 | Rothamsted Research | KWS Siskin | 1 (R) | A02 | I02 | MT856831 |
| Zt_356 | RR17.7 | Western Europe | England | 2017 | Rothamsted Research | KWS Siskin | 1 (R) | A02 | I02 | MT856831 |
| Zt_357 | RR17.8 | Western Europe | England | 2017 | Rothamsted Research | KWS Siskin | 1 (R) | A02 | I02 | MT856831 |
| Zt_358 | T1-A 2017.15 | Western Europe | Ireland | 2017 | Rothamsted Research | KWS Lumos | 7 (S) | A02 | I02 | MT856831 |
| Zt_359 | T1-A 2017.3 | Western Europe | Ireland | 2017 | Rothamsted Research | KWS Lumos | 7 (S) | A02 | I02 | MT856831 |
| Zt_360 | T1-A 2017.43 | Western Europe | Ireland | 2017 | Rothamsted Research | KWS Lumos | 7 (S) | A02 | I02 | MT856831 |
| Zt_361 | T1-A 2017.47 | Western Europe | Ireland | 2017 | Rothamsted Research | KWS Lumos | 7 (S) | A02 | I02 | MT856831 |
| Zt_362 | T1-A 2017.6 | Western Europe | Ireland | 2017 | Rothamsted Research | KWS Lumos | 7 (S) | A02 | I02 | MT856831 |
| Zt_363 | S1-B.15 | Western Europe | Scotland | 2017 | Rothamsted Research | ND | ND | A02 | I02 | MT856831 |
| Zt_364 | S1-B.21 | Western Europe | Scotland | 2017 | Rothamsted Research | ND | ND | A02 | I02 | MT856831 |
| Zt_365 | S1-B.3 | Western Europe | Scotland | 2017 | Rothamsted Research | ND | ND | A02 | I02 | MT856831 |
| Zt_366 | S1-B.4 | Western Europe | Scotland | 2017 | Rothamsted Research | ND | ND | A02 | I02 | MT856831 |
| Zt_367 | S1-B.5 | Western Europe | Scotland | 2017 | Rothamsted Research | ND | ND | A02 | I02 | MT856831 |
| Zt_368 | S.t. DP 0495 | Western Europe | ND | ND | Rothamsted Research | ND | ND | A02 | I02 | MT856831 |
| Zt_369 | M3 2016.17 | Western Europe | England | 2016 | Rothamsted Research | Cordiale | 7 (S) | A02 | I02 | MT856831 |
| Zt_370 | Ztr(ii)2015 | Western Europe | England | 2016 | Rothamsted Research | ND | ND | A16 | I13 | MT856845 |
| Zt_371 | T1-A 2017.24 | Western Europe | Ireland | 2017 | Rothamsted Research | KWS Lumos | 7 (S) | A16 | I13 | MT856845 |
| Zt_372 | 16 | Western Europe | France | 2015 | Rothamsted Research | Trapez | 1 (R) | A07 | I05 | MT856836 |
| Zt_373 | Syn2.12 | Western Europe | France | 2015 | Rothamsted Research | Cellule | ND | A07 | I05 | MT856836 |
| Zt_374 | Syn2.9 | Western Europe | France | 2015 | Rothamsted Research | Cellule | ND | A07 | I05 | MT856836 |
| Zt_375 | 018A | Western Europe | England | 2016 | Rothamsted Research | Solace | 1 (R) | A07 | I05 | MT856836 |
| Zt_376 | 049A | Western Europe | England | 2016 | Rothamsted Research | Amplify | ND | A07 | I05 | MT856836 |
| Zt_377 | 044C | Western Europe | England | 2016 | Rothamsted Research | Stratosphere | ND | A07 | I05 | MT856836 |
| Zt_378 | Syn 2.8 | Western Europe | France | 2015 | Rothamsted Research | Cellule | ND | A41 | I33 | MT856868 |
| Zt_379 | GS26.33 | Western Europe | England | 2015 | Rothamsted Research | Cougar | 1 (R) | A04 | I03 | MT856833 |
| Zt_380 | 4 | Western Europe | England | 2015 | Rothamsted Research | KWS Cashel | 3 (S) | A21 | I17 | MT856849 |
| Zt_381 | 13 | Western Europe | Germany | 2015 | Rothamsted Research | JB Asano | ND | A21 | I17 | MT856849 |

#(S) - *Stb6* haplotype determining susceptibility to *Z. tritici* IPO323 posessing an avirulence isoform of AvrStb6;
(R) - *Stb6* haplotype determining resistance to *Z. tritici* IPO323 posessing an avirulence isoform of AvrStb6;
blue text - *Stb6* haplotype data deterimend in this study;
black text - *Stb6* haplotype data obtained in the previous study by Saintenac *et al.* (2018).

*ND - no data.

**Table S2.** Primers used in this study.

| Primer Name | 5' to 3' Sequence | Purpose |
|---|---|---|
| avrstb6.f1 | CACTTCTTTCCACAACTCCCACTT | Amplification and sequencing of *AvrStb6* |
| avrstb6.f3 | ATCAACTTCCTCTCAACCAAGACC | |
| avrstb6.r1 | CCTACATTGGCAGCATCAAAATCA | |
| 8311F19 | CGCGGTTCCAGTCACATCAC | Amplification and sequencing of *Stb6* |
| 8311F3 | CCGTTTAGCTCGTGTTGTGC | |
| 8311R5F | CTGGACCGCTGGACTTCGAG | |
| 1186R2 | GAGCAAGCTTTCAATTACAGGAG | |
| 13609F1 | CTGAAAAAAAATACGAGGCCATGA | |
| 8311F16 | GCGACATGGTAGCTCAATCAAA | |
| 8311R16 | TTCCTTCCATGGTCGGTAACTT | |
| JPG G6PDH F1 | GCGGCTACTTTGACGAGTTC | RT-qPCR analysis of *AvrStb6* expression |
| JPG G6PDH R1 | GATCCGTCAAGCGACTTCTC | |
| AvrStb6 F6b | TTCTACAAGGCTTCCTCGC | |
| AvrStb6 R6b | GCTTTCCGTCGTGGCAGAA | |
